# *DOMINANT AWN INHIBITOR* encodes the ALOG protein originating from gene duplication and inhibits awn elongation by suppressing cell proliferation and elongation in sorghum

**DOI:** 10.1101/2021.09.29.462495

**Authors:** Hideki Takanashi, Hiromi Kajiya-Kanegae, Asuka Nishimura, Junko Yamada, Motoyuki Ishimori, Masaaki Kobayashi, Kentaro Yano, Hiroyoshi Iwata, Nobuhiro Tsutsumi, Wataru Sakamoto

## Abstract

The awn, a needle-like structure extending from the tip of the lemma in grass species, plays a role in environmental adaptation and fitness. In some crops, awns appear to have been eliminated during domestication. Although numerous genes involved in awn development have been identified, several dominant genes that eliminate awns are also known to exist. For example, in sorghum (*Sorghum bicolor*), the dominant awn inhibiting gene has been known since 1921; however, its molecular features remain uncharacterized. In this study, we conducted quantitative trait locus analysis and a genome-wide association study of awn-related traits in sorghum and identified *DOMINANT AWN INHIBITOR* (*DAI*), which encodes the ALOG family protein on chromosome 3. *DAI* appeared to be present in most awnless sorghum cultivars, likely because of its effectiveness. Detailed analysis of the ALOG protein family in cereals revealed that *DAI* originated from duplication of its twin paralog (*DAI*^ori^) on chromosome 10. Observations of immature awns in near-isogenic lines revealed that DAI inhibits awn elongation by suppressing both cell proliferation and elongation. We also found that only *DAI* gained a novel function to inhibit awn elongation through an awn-specific expression pattern distinct from that of *DAI*^ori^. Interestingly, heterologous expression of *DAI* with its own promoter in rice inhibited awn elongation in the awned cultivar Kasalath. We found that *DAI* originated from gene duplication, providing an interesting example of gain-of-function that occurs only in sorghum but shares its functionality with rice and sorghum.

## Introduction

In grass species, the awn is a needle-like structure that forms at the tip of the lemma. Awns, with a bristle-like architecture, play numerous important roles such as preventing the consumption of grains by animals (Grundbacher 1963, Hua et al. 2015), helping seeds burrow into the soil (Elbaum et al. 2007), and promoting seed dispersal (Sato et al. 1996, Cavanagh et al. 2020). Moreover, awns significantly contribute to photosynthesis and grain yield, particularly in crops such as barley (Kjack and Witters 1974) and wheat (Motzo and Giunta 2002, Li et al. 2006). Wild species of cereal crops tend to have long awns, and numerous genetic studies have demonstrated that multiple factors regulate awn formation and elongation. Although important in wild species, awns in cereal crops often disturb modern agriculture by hindering manual harvesting and reducing the feed value of straw for livestock (Takahashi 1955). Thus, awns were partially or completely eliminated during domestication (Takahashi 1955, Tatsumi and Kawano 1972).

Extensive studies of awn formation and elongation in rice revealed that awnless cultivars were established by sequentially selecting loss-of-function alleles or genes involved in awn formation and elongation during domestication (Amarasinghe et al. 2020, Bessho-Uehara et al. 2021). *An-1* [also known as *REGULATOR OF AWN ELONGATION 1* (*RAE1*); hereinafter referred to as *An-1/RAE1*] encodes a basic helix-loop-helix transcription factor expressed at the apex of the lemma primordia, which causes continuous cell division for awn elongation (Luo et al. 2013, Furuta et al. 2015). LONG and BARBED AWN1 (*LABA1*)/*An-2* encodes a cytokinin-activating enzyme that positively regulates awn elongation (Gu et al. 2015, Hua et al. 2015). *RAE2/GAD1/GLA* encodes Epidermal Patterning Factor-Like protein 1, which acts as a small secretory peptide in the panicles (Bessho-Uehara et al. 2016, Jin et al. 2016, Yano et al. 2016, Zhang et al. 2019). Two other genes, DROOPING LEAF (*DL*), a member of the *YABBY* gene family, and *OsETTIN2* (*OsETT2*) which encodes an auxin-responsive factor also promote awn elongation in rice (Toriba and Hirano 2014); however, these genes are unlikely to be involved in domestication. In barley, short awn 2 (*lks2, for length 2*), which encodes a SHORT INTERNODES family transcription factor, regulates the awn length and pistil morphology (Taketa et al. 2011, Yuo et al. 2012). Genes involved in brassinosteroid biosynthesis or signaling have also been shown to pleiotropically affect awn length (Dockter et al. 2014).

In contrast to these positive factors affecting awn development, inhibitory factors that diminish the awn are also known to exist, including Hooded (*Hd*), Tipped1 (*B1*), and Tipped2 (*B2*) in wheat (Watkins and Ellerton 1940, Kato et al. 1998, Sourdille et al. 2002, Yoshioka et al. 2017). *B1* was recently shown to encode a C2H2 zinc finger protein with ethylene-responsive element binding factor-associated amphiphilic repression motifs that putatively functions as a transcriptional repressor (DeWitt et al. 2020, Huang et al. 2020). Interestingly, such inhibitory factors have not been reported in rice despite extensive studies. Thus, complex and distinct genetic networks may control awn formation in a species-specific manner.

Sorghum (*Sorghum bicolor*) also displays differences in awn morphology among cultivars; however, unlike the studies described above, the genes responsible for awn development are poorly understood. Sorghum is the fifth-highest produced cereal worldwide (faostat.fao.org) and is a C4 crop with a higher stress tolerance than other major cereals (Tuinstra et al. 1997, Ogbaga et al. 2014). In addition, sorghum is a suitable genetic model for C4 grasses because of its high morphological diversity and relatively small genome size (approximately 732 Mb) compared to those of other C4 grasses (Price et al. 2005, Paterson et al. 2009, McCormick et al. 2018). In our recent study, we used a next-generation sequencing approach, restriction site-associated DNA sequencing (RAD-seq), combined with the use of recombinant inbred lines (RILs) derived from a cross between BTx623 and Japanese Takakibi NOG to establish a high-density genetic map that allowed for the investigation of important traits by detecting quantitative trait loci (QTLs) (Kajiya-Kanegae et al. 2020, Jing et al. 2021, Takanashi et al. 2021, Wahinya et al. 2022). We predicted that awn development can also be investigated as the parents used in the RILs displayed contrasting awn morphologies.

In the present study, we conducted QTL analysis using RILs and genome-wide association studies (GWAS) with sorghum germplasm to examine awn-related traits to improve the understanding of awn regulation in sorghum. The results of our QTL analysis for the length and presence or absence of the awn support that a single gene, *DOMINANT AWN INHIBITOR* (*DAI*) on chromosome 3, inhibits awn elongation. We found that *DAI* encodes a protein belonging to the ALOG protein family, whose function has been characterized in cereal species, possibly as a transcription factor and for negatively regulating cell proliferation and expansion in organ development. Indeed, DAI was shown to inhibit awn elongation by suppressing cell proliferation and elongation. *DAI* likely originated from gene duplication of its paralog on chromosome 10, the role of which remains unclear. Interestingly, heterologous expression of sorghum *DAI* in rice made the awned rice cultivar ‘Kasalath’ awnless. Together, these results suggest that common mechanisms exist to inhibit awn elongation in cereal crops; however, the emergence of such inhibitors differs among species.

## Results

### Awned and awnless: morphologies used in this study

Sorghum has two types of spikelets: sessile spikelets (SSs) directly attached to the inflorescence branch and pedicellate spikelets (PSs) attached to the inflorescence branch via a pedicel. Unlike the PS, which has only incomplete floral organs, the SS has a complete floret consisting of a lemma, palea, two lodicules, three anthers, and a pistil; therefore, only the SS can produce a grain (see Fig. 1 of Takanashi et al. 2021). The awn, a needle-like appendage that forms at the tip of the lemma, is formed only in the SS. Recently, we established and reported a sorghum RIL derived from a parental cross between BTx623 and the Japanese landrace NOG (Kajiya-Kanegae et al. 2020). These parents and RILs showed a large difference in awn length (AL) (Fig. 1A and 1B; BTx623 is awnless, whereas NOG is awned). Therefore, QTL analysis of awn-related traits using the RILs was performed using two generations (F_6_ and F_7_) cultivated in Okayama (F_6_, greenhouse, in 2014) and Tokyo (F_7_, field, in 2015) to validate the reproducibility.

**Figure 1.**
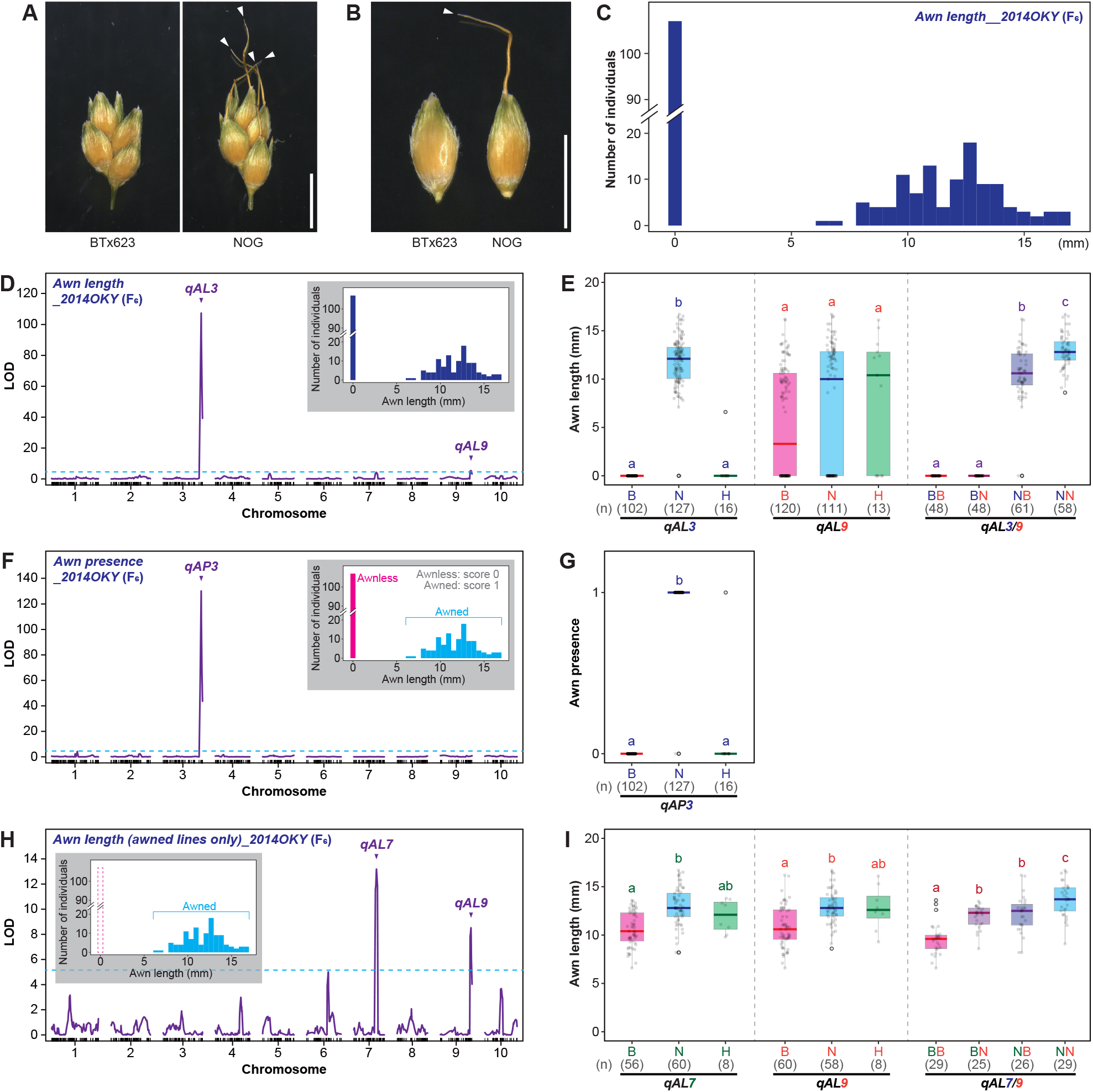
Quantitative trait locus (QTL) analysis of awn-related traits using the sorghum recombinant inbred line (RIL) population derived from BTx623 and NOG. (A) Secondary branches of BTx623 and NOG. (B) Enlarged view of sessile spikelets (SSs) from BTx623 and NOG. Arrowheads indicate the awn tips. (C) Frequency distribution of awn length (AL) in the RIL population (F_6_ generation) cultivated in Okayama in 2014. Results of QTL analysis for awn-related traits (D, F, H) and their allelic effects (E, G, I). (D, E) AL of all lines, (F, G) awn presence (AP), and (H, I) AL of awned lines only. (D, F, H) Logarithm of odds (LOD) profiles obtained by composite interval mapping (CIM). The horizontal dotted lines represent the threshold of the 1,000× permutation test (*P* < 0.05). (E, G, I) Contributions of single-nucleotide polymorphism genotypes of the nearest marker to significant QTLs. Box plots show the effects of the nearest marker genotypes for each QTL or allelic combination of QTLs. B: BTx623-type homogeneous allele; N: NOG-type homogeneous allele; H: heterogeneous. Number of lines corresponding to each genotype is shown in parentheses. Different letters denote significant differences according to the Tukey–Kramer test (*P* < 0.01). Scale bars = 5 mm in (A) and (B).

### QTL analysis of awn-related traits identified a dominant allele that inhibits awn formation or elongation

Measurement of the AL in 245 RIL F_6_ individuals revealed that approximately half of the RILs were awnless, like BTx623 (Fig. 1C). We next performed composite interval mapping of the QTL using AL data of RILs with 1,789 single-nucleotide polymorphism (SNP) markers determined using RAD-seq (Kajiya-Kanegae et al. 2020). We detected a remarkable QTL on chromosome 3, *qAL3* (Table 1). An additional QTL peak was found on chromosome 9 as *qAL9* (Fig. 1D, Table 1). For *qAL3*, the effect of increased AL was attributed to the NOG allele (Fig. 1E, left). Notably, all RIL individuals homozygous for the BTx623 allele or heterozygous for *qAL3* were awnless, indicating that *qAL3* inhibits awn formation or elongation in a dominant manner. Consistent with this observation, the F_1_ plants had no awns in the SSs (Supplementary Fig. 1A). The effect of *qAL9* on AL was unclear, likely because of the superior effect of *qAL3* (Fig. 1E, middle). Nevertheless, the NOG allele for *qAL9* appeared to increase the AL when the allele of *qAL3* was limited to the NOG type (Fig. 1E, right).

**Table 1.** Summary of awn-related quantitative trait loci (QTLs) detected in this study.

| Trait | Year_location | Chr | QTL ID | Position (cM) | Nearest marker | Marker interval | LOD | PVE (%) | AE |
| --- | --- | --- | --- | --- | --- | --- | --- | --- | --- |
| Awn length (AL) | 2014_OKY | 3 | *qAL3* | 76.62 | Chr03:72655415 | Chr03:71923025–72847396 | 116.4 | 90.6 | 5.75 |
|  | 2015_TKY | 3 | *qAL3* | 76.62 | Chr03:72655415 | Chr03:71923025–72847396 | 100.8 | 88.7 | 4.58 |
|  | 2014_OKY | 9 | *qAL9* | 59.76 | Chr09:57681027 | Chr09:56467317–58331590 | 7.0 | 1.3 | 0.64 |
| Awn presence (AP) | 2014_OKY | 3 | *qAP3* | 76.62 | Chr03:72655415 | Chr03:71923025–72847396 | 179.1 | 97.8 | 0.49 |
|  | 2015_TKY | 3 | *qAP3* | 76.62 | Chr03:72655415 | Chr03:71923025–72847396 | 120.8 | 92.7 | 0.48 |
| Awn length (awned lines only) | 2014_OKY | 7 | *qAL7* | 47.27 | Chr07:59733246 | Chr07:58230302–60603839 | 8.3 | 23.8 | 1.06 |
|  | 2015_TKY | 7 | *qAL7* | 47.40 | Chr07:59733246 | Chr07:58230302–60603839 | 3.5 | 10.7 | 0.64 |
|  | 2014_OKY | 9 | *qAL9* | 59.76 | Chr09:57681027 | Chr09:56467317–57917278 | 5.9 | 16.2 | 0.87 |
|  | 2015_TKY | 9 | *qAL9* | 60.06 | Chr09:57681027 | Chr09:56467317–58331590 | 4.8 | 15.0 | 0.76 |
| OKY, Okayama; TKY, Tokyo. Marker intervals were estimated based on confidence intervals (2.0-LOD). PVE: Phenotypic variance explained. For the additive effects (AE), positive values indicate that alleles from NOG increased the trait score. |  |  |  |  |  |  |  |  |  |

To evaluate whether *qAL3* determines the awn presence (AP) rather than the AL, we conducted QTL analysis using data converted from AL data to AP (re-scored as awnless = 0 and awned = 1). We again detected a single QTL on chromosome 3 (*qAP3*) in the same position as *qAL3* but with a more intense effect (Fig. 1F, Table 1). The allelic effect of *qAP3* was similar to that of *qAL3;* the BTx623 allele predominantly led to an awnless phenotype (Fig. 1G). Together, these results suggest that the single locus corresponding to *qAL3/qAP3* is the dominant inhibitor of awn formation or elongation.

### Existence of two QTLs controlling awn length

We hypothesized that minor QTLs were not detectable because the strong awn inhibitory effect of *qAL3/qAP3* obscured the effect of minor QTLs. To further investigate the control of AL, we performed QTL analysis using AL data from awned lines only that were not affected by the awn inhibitory effect of *qAL3/qAP3*. As expected, we detected two additional QTLs, *qAL7* on chromosome 7 and *qAL9* on chromosome 9 (Fig. 1H, Table 1). Considering that the nearest marker and marker interval of *qAL9* detected here was similar to that of *qAL9* in the initial QTL analysis that included awnless individuals, both were considered to represent the same QTL. The increased AL was attributed to NOG alleles for both *qAL7* and *qAL9* (Fig. 1I). The QTLs (*qAL3/qAP3, qAL7*, and *qAL9*) were also detected in the QTL analysis using the datasets of 2015 cultivation in Tokyo, indicating high reproducibility for all QTLs (Supplementary Fig. 1B–H, Table 1). Based on these findings, the dominant inhibitory effect of *qAL3/qAP3* was apparent; thus, we further characterized *qAL3/qAP3* hereafter.

### GWAS revealed the widespread distribution of qAL3/qAP3 among sorghum cultivars

To clarify whether the inhibitory effect of *qAL3/qAP3* is globally involved in awn regulation in sorghum, we conducted GWASs for AL using 289 accessions of sorghum germplasm, whose data were collected during cultivation in Tokyo for three years (2015, 2019, and 2020). Our germplasm collection exhibited large variability in the AL (Fig. 2A), and more than half of the accessions were awnless (Fig. 2B). First, an AL GWAS showed significant associations on chromosome 3, with a peak at Chr03:72641259 (Fig. 2C). This region was close to *qAL3*/*qAP3*, and the GWAS peak and nearest marker of *qAL3*/*qAP3* were only 14 kb apart (Fig. 2D). The effect of the peak genotype was the same as that of *qAL3/qAP3* (Fig. 2E). Subsequently, we performed an AP GWAS and detected similar associations with those in AL on chromosome 3 (Fig. 2C insert). These detected associations were also found in GWAS using datasets from the 2019 and 2020 cultivations, indicating the high reproducibility of this association (Supplementary Fig. 1I and 1J). Taken together, we considered that *qAL3*/*qAP3* detected in QTL analysis and the GWAS peaks are likely to represent the same gene.

**Figure 2.**
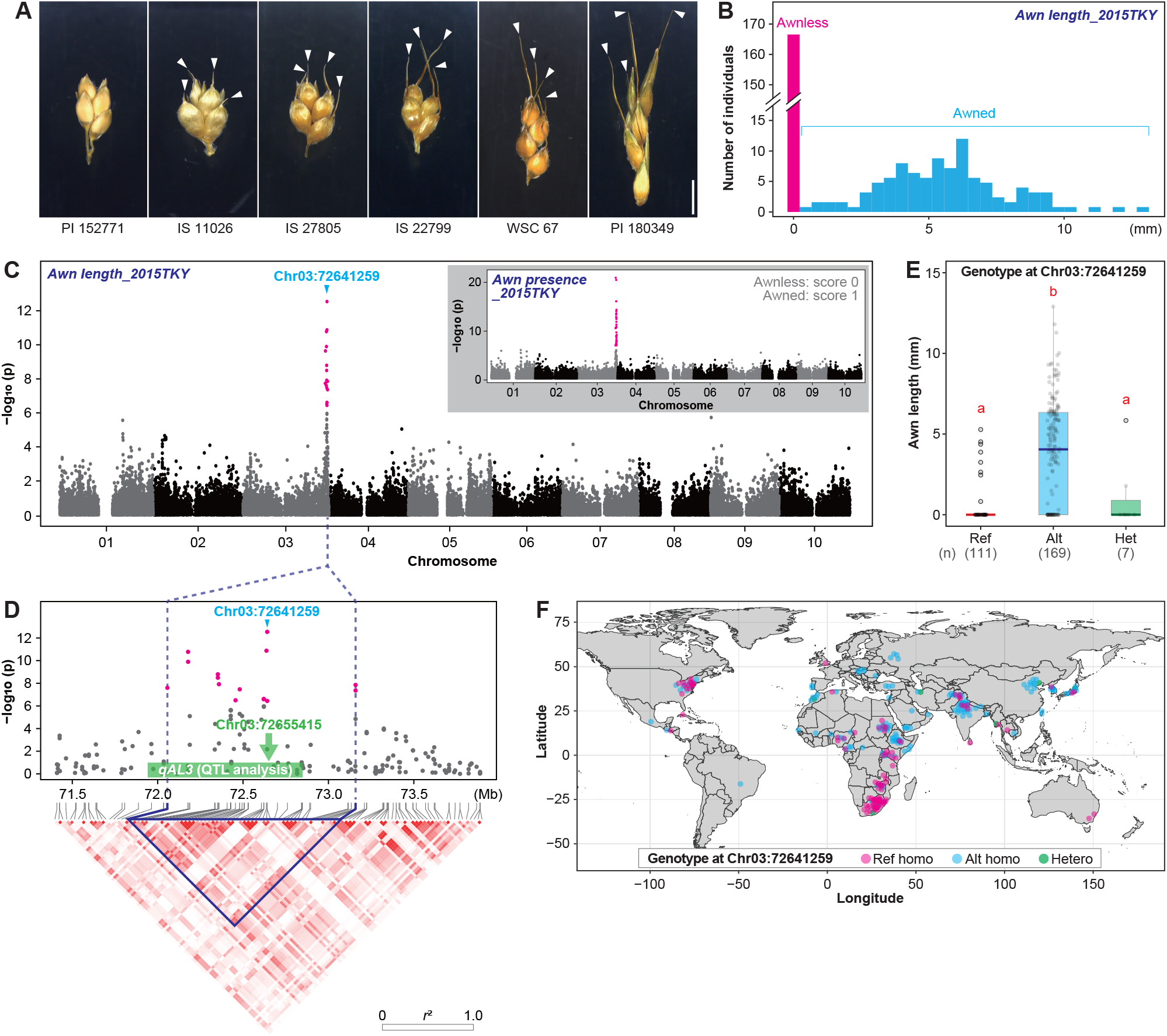
Genome-wide association study for awn-related traits using the sorghum germplasm. (A) Examples of secondary branches from the panicles of sorghum germplasm. Accession ID is shown at the bottom of each panel. Arrowheads indicate awn tips. (B) Frequency distribution of awn length (AL) in the sorghum germplasm cultivated at Tokyo in 2015. (C) Manhattan plot for AL and awn presence (AP) (insert). Dots in magenta indicate the single-nucleotide polymorphisms (SNPs) significantly associated with traits (Benjamini–Hochberg-corrected significance, *P* < 0.001). (D) Local Manhattan plot (top) and linkage disequilibrium heatmap (bottom) surrounding the peak on chromosome 3. A bar and arrow in green indicate the marker interval and position of the nearest marker of *qAL3/qAP3* in QTL analysis, respectively. (E) Box plot showing the effects of the peak SNP (Chr03:72641259) genotype for AL. Ref, reference-type homogeneous allele; Alt, altered-type homogeneous allele; H, heterogeneous. Number of lines that corresponded to each genotype is shown in parentheses. Different letters denote significant differences according to the Tukey–Kramer test (*P* < 0.01). (F) Geographic distribution of alleles at the peak SNP (Chr03:72641259), color-coded by allele (Ref homo: magenta, Alt homo: cyan, Hetero: green). Scale bar = 5 mm in (A).

To assess the geographic distribution of alleles at the peak SNP detected using GWAS, we color-coded the individual accessions by genotype and plotted the origins of individual accessions on a world map (Fig. 2F). Awnless accessions with a Ref-type allele (BTx623 type) were abundant in South Africa and North America, whereas awned accessions with an Alt-type allele (NOG type) were common in Asia, North Africa, and Europe. However, whether this geographic distribution reflects adaptation to regional climates or resulted from human selection for different uses is unclear.

### Candidate of dominant awn-inhibiting gene

We compared the reference genome sequence of BTx623 with those from NOG, which were determined using our re-sequencing data (accession number: DRA008159). A region of approximately 5 kb containing a single gene, *Sobic003G421300*, was missing in NOG (Supplementary Fig. 2A). Indeed, we verified that a 5.46-kb fragment containing this gene was missing in NOG using genomic PCR and Sanger sequencing (Supplementary Fig. 2B and 2C, Fig. 3A). The peak SNP of the GWAS was approximately 5.7 kb upstream of the transcriptional start site of *Sobic003G421300*. According to the Phytozome database (Goodstein et al. 2012), *Sobic003G421300* encodes a putative transcription factor containing the *Arabidopsis* LSH1 and *Oryza* G1 (ALOG) domain (Fig. 3B). Because an ALOG protein was reported to inhibit lateral organ development by suppressing cell division (Naramoto et al. 2019), we predicted that Sobic003G421300 also inhibits awn formation or elongation, and this gene was named as *DOMINANT AWN INHIBITOR* (*DAI*). Because known ALOG family proteins act as transcription factors, we hypothesized that DAI is a transcription factor that inhibits awn formation or elongation. To test this possibility, we transiently expressed green fluorescent protein (GFP) or DAI-GFP fusion protein driven by the cauliflower mosaic virus *35S* promoter in onion epidermal cells by particle bombardment. Although control GFP was localized in the cytoplasm and nucleus (Fig. 3C upper), the DAI-GFP fusion protein was exclusively localized in the nucleus (Fig. 3C bottom), suggesting that DAI is a transcription factor that negatively regulates awn formation or elongation.

**Figure 3.**
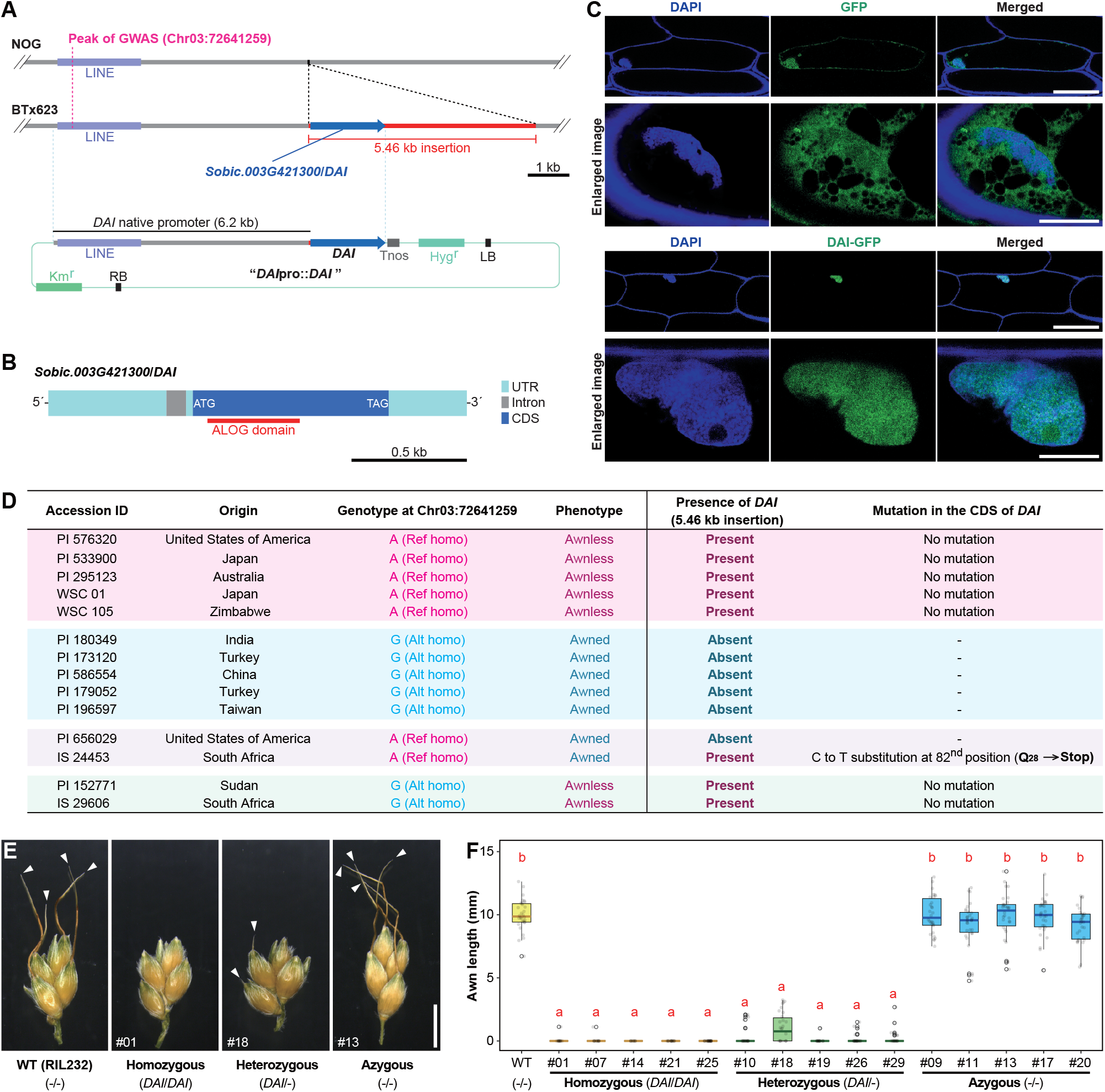
Confirmation of the *DAI* locus encoding the ALOG protein responsible for awn inhibition. (A) Schematic representation of the genomic region surrounding *DAI* on chromosome 3 in the NOG and BTx623 genome (upper), and the *DAI*pro::*DAI* plasmid used for complementation analysis (bottom). (B) Predicted gene structure of *DAI* in BTx623. Pale blue, gray, and blue boxes indicate untranslated region (UTR), intron, and coding sequence (CDS), respectively. A red line indicates a region coding for the ALOG domain. (C) Localization of green florescent protein (GFP) (upper) and DAI-GFP fusion protein (bottom) in onion epidermal cells. (D) Relationship among genotype of the peak single-nucleotide polymorphism (SNP) in genome-wide association study (Chr03:72641259), awn phenotype, presence/absence of *DAI*, and mutations within the CDS of *DAI* in 14 accessions of sorghum germplasm. (E) Representative images of secondary branches from panicles of the awned parental line (RIL232), homozygous transgenic plant (*DAI/DAI*), heterozygous transgenic plant (*DAI*/-), and azygous plant (-/-). Arrowheads indicate awn tips. (F) Box plot showing AL of the RIL232, homozygous, heterozygous, and azygous plants. Different letters denote significant differences according to the Tukey– Kramer test (*P* < 0.01). Scale bars = 100 μm in (C, 10 μm in enlarged images), 5 mm in (E).

### Confirmation by genotyping and complementation analysis

Additional experiments were conducted to verify that *DAI* corresponded to *qAL3/qAP3*. First, we examined the link between the awn phenotype and presence of *DAI*. Although the peak SNP correlated well with the awn phenotypes in most accessions (Fig. 2E), a few accessions showed discrepant phenotypes, including the awned with the Ref-type (BTx623) allele and awnless with the Alt-type (NOG) allele. Therefore, we selected 14 accessions, including those showing contrasting genotypes (PI 656029, IS 24453, PI 152771, and IS 29606; Fig. 3D, bottom) and examined whether they had a 5.46-kb fragment containing *DAI* (*DAI* fragment) and/or whether they had additional mutations in *DAI* using Sanger sequencing (Fig. 3D). As expected, the peak genotype and presence/absence of the *DAI* fragment matched for ten accessions whose awn phenotypes were consistent with the peak genotype. For the remaining four accessions showing this discrepancy, our experimental reassessment clarified that three accessions (PI 656029, PI 152771, and IS 29606) were consistent between the awn phenotype and presence/absence of the *DAI* fragment. In contrast, IS 24453 was remarkably different because it was awned despite the presence of the *DAI* fragment. However, Sanger sequencing of IS 24453 revealed that a single-nucleotide substitution occurred in *DAI* (82 bp from the translation start point; C to T), resulting in a premature stop codon that truncated DAI to only 28 amino acids (282 amino acids in wildtype). Thus, IS24453 represents a novel haplotype. These results suggest that *DAI* determines the absence of awns in sorghum.

To determine whether *DAI* prevents the appearance of awn, we performed complementation analysis in sorghum using *Agrobacterium*-mediated transformation with tissue cultures derived from immature embryos. We cloned the genomic fragment of BTx623 containing the putative promoter region (approximately 6.2 kb) and *DAI* into a binary vector (*DAI*pro::*DAI*) (Fig. 3A, bottom). RIL232 is an awned line in our RIL population and was used as the recipient. The obtained transformant (T_0_) showed a single insertion based on the segregation ratio of the T_1_ progeny. Genotyping of the T_1_ plants was performed to distinguish homozygous (*DAI/DAI*), heterozygous (*DAI*/-), or azygous (-/-) individuals using quantitative PCR. Five T_1_ plants of each genotype were selected for SS observation and AL measurement. We found that homozygous plants completely lost their awns, heterozygous plants were partially awned but most had lost their awns, and azygous plants had the same level of awns as the parent (Fig. 3E and 3F). Together, these results demonstrate that *DAI* is responsible for *qAL3*/*qAP3*.

### DAI originated from a gene duplication

*DAI* is present in the 5.64-kb genomic fragment missing in accessions such as NOG. We next investigated the origin of this fragment; unexpectedly, the BLASTP search revealed the presence of one protein encoded by *Sobic.010G223100* on chromosome 10, which showed 100% amino acid sequence identity with DAI. Comparison of the coding sequences (CDSs) revealed only one synonymous substitution at nucleotide 603 bp downstream of the translation start point. Moreover, *DAI* and *Sobic.010G223100* were similar not only between CDSs but also in the regions encompassing 5.15 kb (Supplementary Fig. 3). Considering that *Sobic.010G223100* is also present in both BTx623 and NOG and that the awned phenotype predates the awnless phenotype during varietal differentiation, the 5.46-kb fragment on chromosome 3 likely resulted from a duplication of *Sobic.010G223100* in the BTx623 ancestor, rather than from a deletion in the NOG ancestor. Therefore, we named this *DAI* homolog on chromosome 10 as *DAI*^ori^. To further verify this hypothesis, synteny analysis was performed to compare the corresponding chromosomal regions among three other grass species [rice (Os), *Brachypodium* (Bd), and maize (Zm) in Fig. 4A]. We detected syntenic blocks that corresponded to the region around *DAI* on sorghum chromosome 3, Os chromosome 1, Bd chromosome 2, and Zm chromosomes 3 and 8. However, none of these blocks contained *DAI* homologs in Os, Bd, and Zm (Fig. 4A and 4B upper). In contrast, *DAI*^ori^ orthologs were found in the syntenic blocks of all these grass species (in Os chromosome 6, Bd chromosome 1, and Zm chromosomes 6 and 9) (Fig. 4A and 4B bottom). Only *DAI*^ori^ was conserved in these grass species, whereas the presence of *DAI* correlated well with awnless sorghum accessions. These observations support the hypothesis that *DAI*^ori^ is the origin of *DAI* and its copy may have inserted into chromosome 3 in the BTx623 ancestor.

**Figure 4.**
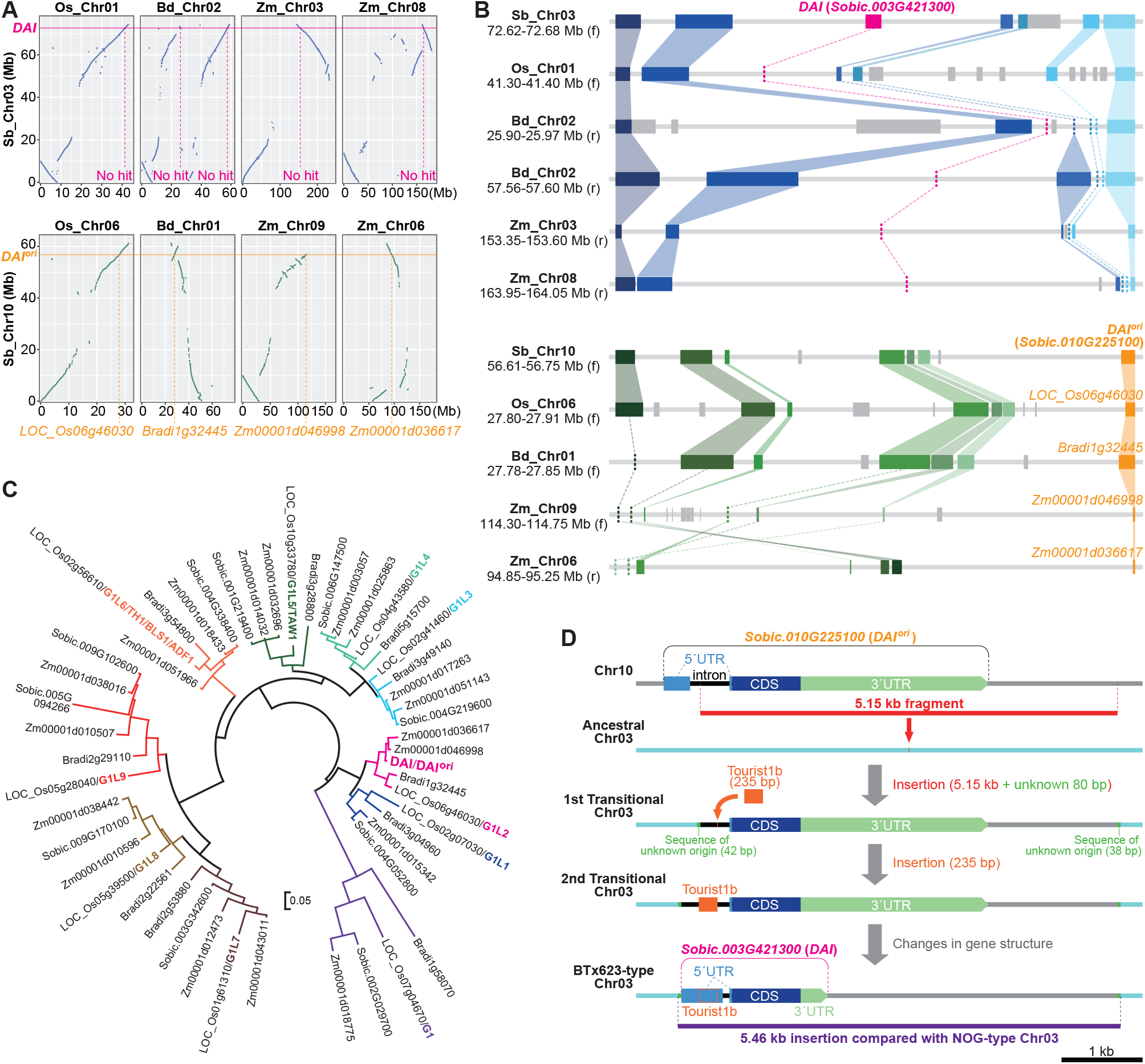
Generation and evolution of *DAI* in sorghum. (A, B) Synteny analysis for *DAI* and its paralogous gene, *DAI*^ori^ among four grass species. (A) Chromosome-level synteny analysis of sorghum chromosome 3 encoding *DAI*, and 10 encoding *DAI*^ori^. Sb: *Sorghum bicolor*, Os: *Oryza sativa*, Bd: *Brachypodium distachyon*, and Zm: *Zea mays*. (B) Schematic diagram of the genomic structures around *DAI*, *DAI*^ori^, and their syntenic regions in four grass species. Boxes in the same color indicate the genomic region containing orthologous genes. Genes shown in dotted lines indicate that the corresponding genes do not exist in this region. ‘f’ and ‘r’ indicate forward and reverse strands, respectively. (C) Phylogenetic tree for DAI/DAI^ori^ and related proteins from the four grass species. (D) Model for the generation and evolution of *DAI*. Gene structures of *DAI* and *DAI*^ori^ are based on the information of the reference genome of *S. bicolor* (v3.1.1).

Next, we performed BLASTP analysis of four grass species using the DAI/DAI^ori^ protein as a query and constructed a phylogenetic tree using the selected proteins. All proteins were grouped into clades represented by each rice G1 family protein (G1, G1L1–L9), and gene duplication was particularly prominent in maize (Fig. 4C). These ALOG family proteins were well-conserved, at least among the four species evaluated. DAI/DAI^ori^ was found to belong to the G1L2 clade of unknown function, which differed from the TAW1 (G1L5 clade), TH1/BLS1/ADF1 (G1L6 clade), and G1 clades, whose functions have been reported (Yoshida et al. 2009, Li et al. 2012, Ma et al. 2013, Yoshida et al. 2013, Sato et al. 2014, Ren et al. 2016, Peng et al. 2017).

We predicted that *DAI* was generated by gene duplication through genome rearrangement (Supplementary Fig. 3 and Fig. 4D). First, a fragment of approximately 5.15 kb encompassing the intron and CDS of *DAI*^ori^ on chromosome 10 appeared to have been duplicated and inserted into chromosome 3 (at approximately 72.6 Mb), followed by addition of an unidentified sequence (total of 80 bp) at both ends; the origin of this sequence is currently unclear because BLAST searches against all species revealed no hits other than its own. Subsequent insertion of a transposable element, Tourist1b, into the intron region may have led to further rearrangement in the untranslated regions and introns, resulting in formation of *DAI* in its present form.

### DAI inhibits awn elongation by suppressing both cell proliferation and elongation

To determine how DAI inhibits awn formation or elongation, we conducted functional analysis using established near-isogenic lines (NILs). Several RILs were heterozygous for *DAI* in the F_6_ population. Of these, we selected an awnless heterozygous line, RIL205, and followed the segregation of *DAI* in subsequent selfed generations (Fig. 5A). Observations of the SSs of the NILs (F_9_ generation) showed that the NIL with the homogeneous BTx623 allele (NIL_B, *DAI/DAI*) lost its awns, whereas the other NIL with the homogeneous NOG allele (NIL_N, -/-) had awns (Fig. 5B). Analysis of immature SSs in NIL_N indicated that awns appeared when the length of the SS reached approximately 2.0 mm (between stages 5 and 6; Fig. 5C). Dissection of immature SSs revealed no apparent differences between NIL_B and NIL_N at stage 5. However, at stage 7, the awn emerged only in NIL_N. Importantly, we observed small awns inside the SSs, even in NIL_B, at both stages (Fig. 5D), suggesting that DAI inhibits awn elongation rather than awn formation. We defined the “dissected awn length (DAL)” as the awn length from the notch of the lemma to the tip (Fig. 5D, DAL) and measured these lengths at stages 5 and 7). The results showed that DAL increased significantly as the developmental stage progressed in NIL_N (from 1.23 to 3.97 mm on an average) but not in NIL_B (from 0.44 to 0.62 mm on an average) (Fig. 5F). In addition, there was a significant difference in the DAL between NIL_B and NIL_N at stage 5. These results confirm that DAI functions from before stage 5 to inhibit awn elongation.

**Figure 5.**
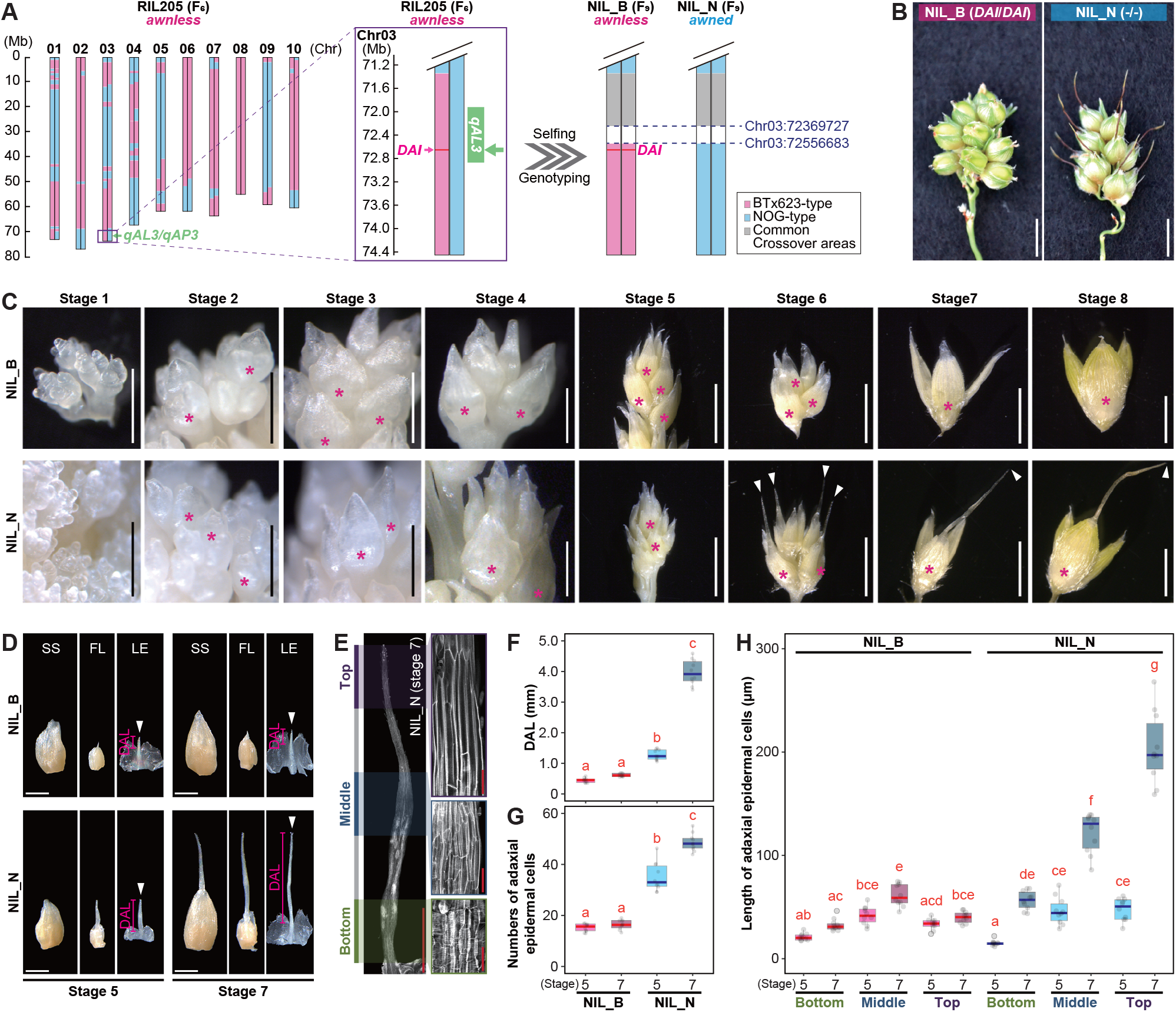
Analysis of how DAI inhibits awn elongation using near-isogenic lines (NILs). (A) Graphical genotypes of RIL205 (F_6_, awnless), one of the heterozygous lines at the genotype of the marker associated with *qAL3/qAP3* (left), and magnified view of the Chr03 terminus for RIL205, NIL with the homogeneous BTx623 allele (NIL_B, F_9_, awnless), and that with the homogeneous NOG allele (NIL_N, F_9_, awned) at *qAL3/qAP3* (right). Magenta and cyan boxes indicate genomic regions from BTx623 and NOG, respectively. PCR genotyping using insertion-deletion polymorphisms revealed that recombination occurred between Chr03:72369727–Chr03:72556683 in both NILs; differences between NILs at the Chr03 terminus were approximately 1.83–2.02 Mb. “Common” indicates that the genotypes (homogeneous allele) are common between both NILs; however, we could not determine whether they were the BTx623-type or NOG-type using PCR genotyping. A bar and arrow in green indicate the marker interval and position of the nearest marker of *qAL3*/*qAP3* in the QTL analysis, respectively. (B, C) Observations of sessile spikelets (SSs) of NILs at various growth stages. (B) Enlarged images of the primary branches at the grain-filling period. Scale bars = 5 mm. (C) Immature SSs of young panicles. Staging of SSs was carried out based on the length of SSs (asterisks). Stage 1: primordial stage, stage 2: 0.3–0.4 mm, stage 3: 0.4–0.6 mm, stage 4: 0.6–0.8 mm, stage 5: 1.5–2.0 mm, stage 6: 2.0–2.5 mm, stage 7: 2.6–3.3 mm, and stage 8: 3.7–4.2 mm, respectively. Scale bars = 0.5 mm in stage 1–4, 2.0 mm in stage 5–8. (D) Step-by-step dissection of immature SSs at stages 5 and 7. See Fig. 6A for details on the components of the SS. FL: floret, LE: lemma (with awn), DAL: dissected awn length. Scale bars = 1.0 mm. Arrowheads in (C) and (D) indicate awn tips. (E) Examples of the dissected awn (NIL_N, stage 7) whose cell walls were visualized using fluorescent staining (calcofluor white). Scale bars = 1.0 mm in the left panel, 50 μm in the right panels. (F–H) Box plots show the DAL (F), numbers of adaxial epidermal cells (G), and lengths of adaxial epidermal cells of the three regions (bottom, middle, and top; see Fig. 5E) (H) for the dissected awns of SSs at stages 5 and 7. Different letters denote significant differences according to the Tukey–Kramer test (*P* < 0.01).

Next, we visualized the cell walls of the dissected awns using fluorescent staining (Fig. 5E) and counted the number of epidermal cells lined up vertically to form the awn at stages 5 and 7. The results showed that the number of adaxial epidermal cells increased significantly in NIL_N (from 35.2 to 48.7 on an average) but did not increase significantly in NIL_B (from 15.3 to 16.4 on an average) (Fig. 5G). We also found a significant difference in cell numbers between NIL_B and NIL_N at stage 5. These results indicate that DAI represses cell proliferation of immature awns. To assess the effect of DAI on cell elongation, we measured the length of epidermal cells in the dissected awns in three different regions (bottom, middle, and top in Fig. 5E). The results indicated that the epidermal cell lengths increased significantly in all regions as the developmental stage progressed in NIL_N but not in NIL_B (Fig. 5H). Taken together, DAI inhibits awn elongation by suppressing both cell proliferation and elongation.

### Spatiotemporal expression of DAI and DAI^ori^ confirms awn-specific expression of DAI

Considering that *DAI* originated from gene duplication of *DAI*^ori^ and that both genes encode identical ALOG proteins, it remains unclear how only *DAI* exhibited a novel function to inhibit awn elongation. In other words, the question is whether the acquisition of the novel function is due to merely increased expression levels through gene duplication or altered expression patterns by obtaining a new promoter. To verify this, we quantified expression levels of *DAI* and *DAI*^ori^ using quantitative reverse transcription (qRT)-PCR. Although almost identical sequences of the two genes limited our spatiotemporal expression analysis, we generated a primer set for qRT-PCR, by which *DAI*-specific expression could be quantified successfully (see Materials and Methods). First, we measured the transcript levels of the two genes in the immature primary branches of NIL_B for each stage (Fig. 5C), as well as in the leaves and roots. The results showed that *DAI*^ori^ expression was higher than that of *DAI*, whereas *DAI* expression was significantly elevated between stages 4 and 5 and was distinct from the *DAI*^ori^ expression pattern (Fig. 6C). Neither of the transcripts accumulated in the leaves, and only *DAI^ori^* was high in the roots.

**Figure 6.**
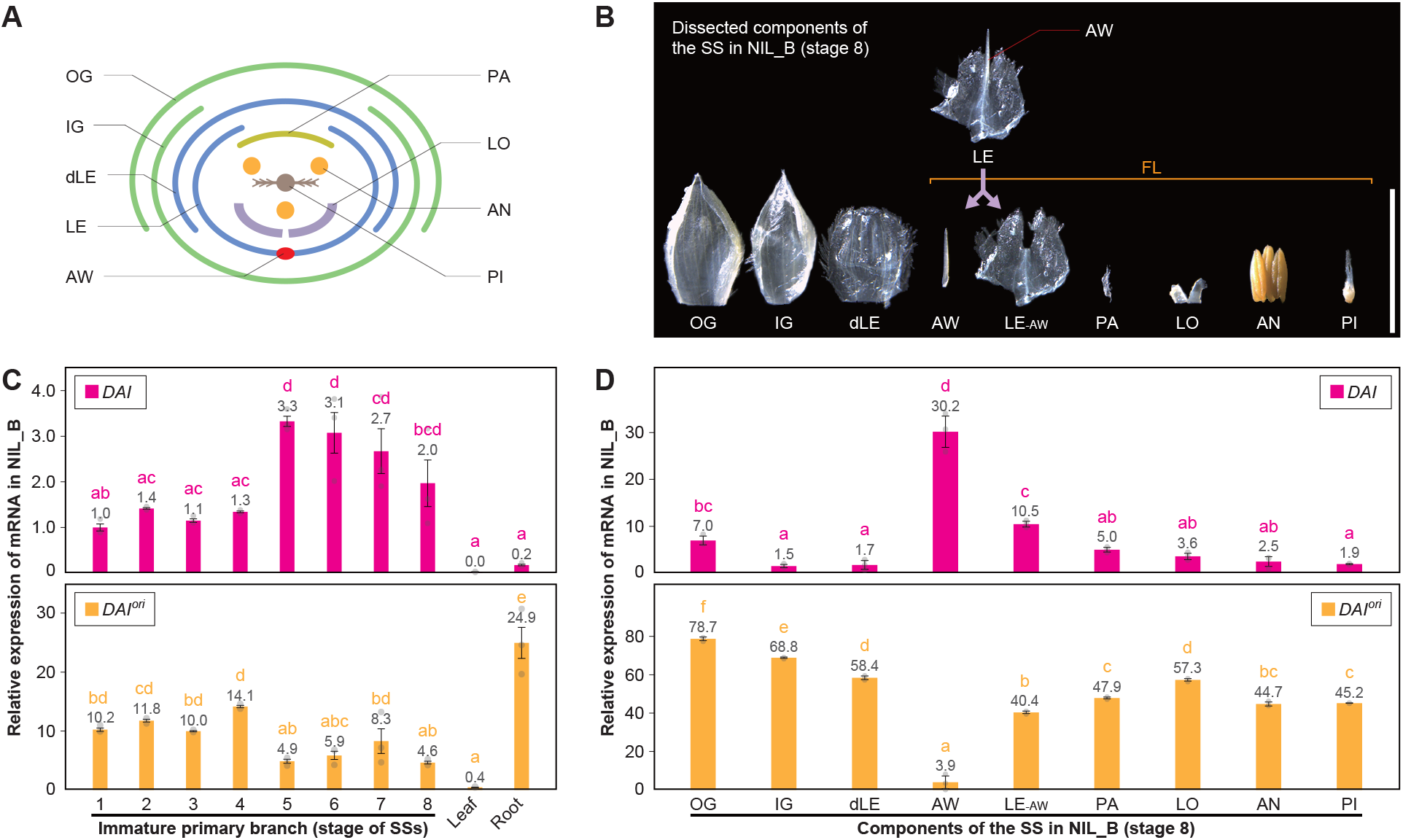
Expression analysis for *DAI* and *DAI*^ori^ in NIL_B using qRT-PCR. (A) Schematic diagram of the internal structure of the sessile spikelets (SS) in sorghum. OG: outer glume, IG: inner glume, dLE: lemma of a degenerated floret, LE: lemma with awn (AW), PA: palea, LO: lodicule, AN: anther, and PI: pistil. (B) Example of the dissected components of the SS in NIL_B at stage 8. LE was separated into two parts: AW and LE without AW (LE-AW). Scale bar = 5 mm. (C, D) Expression patterns of *DAI* and *DAI*^ori^ in immature primary branches, leaf, and root (C), and components of the SS at stage 8 (D) in NIL_B. Error bars indicate standard errors. Different letters denote significant differences according to the Tukey–Kramer test (*P* < 0.01).

To gain insights into the spatiotemporal expression of *DAI* and *DAI*^ori^ in the SS, we dissected all components of the SS at stage 8 and individually quantified the transcript levels of the two genes. In sorghum, each component of a single SS was arranged in the following order: an outer glume, inner glume, lemma of a degenerated floret, lemma with awn (LE), palea, two lodicules, three anthers, and a pistil (Fig. 6A). We separated LE into two parts: awn (AW) and lemma without awn (LE-AW) (Fig. 6B). Our qRT-PCR results revealed that although *DAI*^ori^ was expressed at a higher level than *DAI* in most components, *DAI* exhibited specific accumulation in the awns. In contrast, *DAI*^ori^ was rarely detected in the awns (Fig. 6D). These results suggest that the novel function of DAI in inhibiting awn elongation is by a distinct awn-specific expression pattern rather than by increased expression levels following gene duplication.

### SbDAI can also function as an awn inhibitor in rice

*DAI*, which has acquired a promoter leading to expression in the awn, encodes a conserved ALOG family protein among grass species, suggesting that the whole *DAI* allele, including its own promoter, acts as an awn inhibitor in species other than sorghum. To examine this possibility, we generated transformants using the awned rice cultivar Kasalath as a recipient heterologously expressing *DAI* and observed their spikelets. Using the pHGWFS7 vector as a backbone, the *SbDAI* construct harboring the genomic region of BTx623 (Fig. 7A) was prepared. As a negative control, the corresponding region from NOG was cloned into the gNOG construct (Fig. 7A). Each binary vector carrying *SbDAI* or gNOG was transformed into the awned rice cultivar Kasalth. Individuals with gNOG exhibited an awned phenotype similar to that of Kasalath, whereas those with *SbDAI* showed a completely awnless phenotype (Fig. 7B–D). Three independent transformants (T_0_) obtained for each construct were single-copy transformants, and we determined whether they contained transgene(s) (+ in Fig. 7E) or not (azygous: Az in Fig. 7E) by PCR using DNA isolated from each of the segregated T_1_ populations. A T_1_ plant of each genotype was selected for measuring the AL. The results showed that the awns of individuals with the *SbDAI* transgene were significantly shorter (almost zero) than those of their azygous counterparts (Fig. 7E). These results indicate that *SbDAI* (native promoter + *DAI*) also inhibits awn elongation in heterologous rice plants, supporting the gain-of-function of *DAI* by gene duplication.

**Figure 7.**
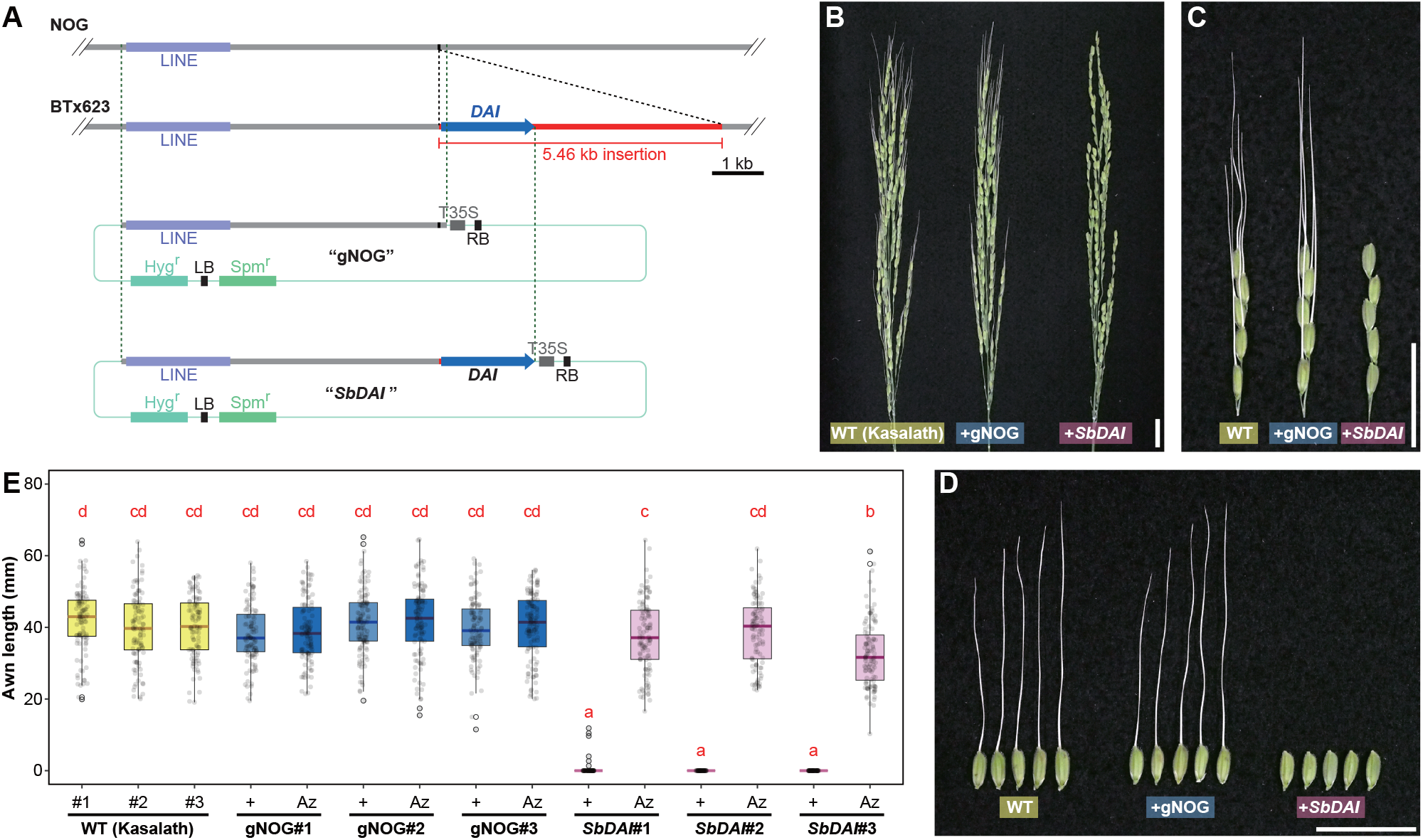
*SbDAI* inhibits awn elongation in the awned rice cultivar Kasalath. (A) Schematics of the gNOG and *SbDAI* plasmids for making transgenic Kasalath. (B–D) Examples of panicles (B), secondary branches (C), and spikelets (D) of wild-type Kasalath, gNOG transgenic line, and *SbDAI* transgenic line. (E) Box plot shows the AL of the wild-type Kasalath, gNOG transgenic lines, *SbDAI* transgenic lines, and their azygous lines. ‘+’ or ‘Az’ indicate the individual has the transgene or does not, respectively. Different letters denote significant differences according to the Tukey–Kramer test (*P* < 0.01). Scale bars = 2 cm in (B–D).

## Discussion

### Identification of DAI as a major gene for the diversification of sorghum awnless cultivars

Awn-related traits in sorghum have been reported previously (Boivin et al. 1999, Tao et al. 2000, Hart et al. 2001) and investigated using GWAS (Shehzad et al. 2009, Zhang et al. 2015, Girma et al. 2019). Notably, the existence of a dominant awn inhibitor locus was documented more than a century ago in sorghum (Vinall and Cron 1921, Sieglinger et al. 1934). These reports underscore the emergence of the awnless trait in the diversification or improvement of sorghum cultivars. However, the molecular identity and cellular dissection of these traits remained unclear until recently. In this study, we focused on the major QTL that regulates awn length and awn presence (*qAL3/AP3*), which was identified in an RIL population derived from BTx623 and NOG (Fig. 1). *qAL3/qAP3* likely maps to the awn trait reported in previous GWASs by Zhang et al. (2015) and Girma et al. (2019).

We previously demonstrated that the combination of BTx623 and NOG is useful for dissecting various traits, such as those related to the photosynthetic capacity during grain filling (Ohnishi et al. 2019), biomass (Kajiya-Kanegae et al. 2020), spikelet morphology (Takanashi et al. 2021), organophosphate pesticide sensitivity (Jing et al. 2021), and seed ionome (Wahinya et al. 2022). Using this population, we cloned *DAI*, which is present only in BTx623 because of gene duplication. Our results, including those of GWAS (Fig. 2), genotyping of other sorghum accessions (Fig. 3), transgenic analyses using sorghum and rice (Fig. 3E, 3F, and 7), and detailed functional analysis of NILs (Fig. 5 and 6), demonstrated that *DAI* inhibits awn elongation in a dominant manner. Moreover, most awnless accessions were accounted for by the presence of functional *DAI* (Fig. 2 and 3), strongly suggesting that *DAI* is critically important for the development of modern awnless sorghum cultivars. *DAI* is useful and easy to genetically manipulate as a dominant factor, for example, in generating awnless F_1_ hybrids. Therefore, *DAI* may have been widely selected during varietal diversification. Whether *DAI* is associated with sorghum domestication remains unclear and should be further examined. Additionally, studies are required to understand the mechanism by which the putative transcription factor DAI inhibits awns by affecting the downstream target genes.

While preparing this manuscript, Zhou et al. (2021) reported a gene named as *awn1*, identical to *DAI* in this study, as a candidate awn repressor in sorghum. Although their study did not provide direct evidence that *awn1/DAI* is the gene responsible for the dominant awn inhibitor, they performed several important analyses, including RNA sequencing and DNA affinity purification sequencing, which led to the identification of genes downstream of *awn1*. The results of their study as well as those of the current study improve the understanding of awn regulation in sorghum.

### DAI is an ALOG family protein and inhibits awn elengation

*DAI* belongs to the ALOG family and differs from any other awn-regulating genes reported in other grass species. In wheat, one of the dominant awn inhibitors, *B1*, encodes a C2H2 zinc finger protein (DeWitt et al. 2020, Huang et al. 2020), suggesting that many types of dominant awn inhibitor genes may exist in grass species. Nevertheless, our heterologous study in rice suggests that the mechanism of suppressing awn elongation manifested by proteins with ALOG domains are shared at least between sorghum and rice (Fig. 7).

ALOG family proteins, which may have emerged during the early evolution of *Streptophyta*, exhibit high sequence conservation among ALOG domains in divergent plant lineages (Naramoto et al. 2020). Among the ALOG family proteins in grass species, G1, TAW1, and TH1/BLS1/AFD1 have been well characterized in rice. LONG STERILE LEMMA1 (G1) is thought to be involved in transcriptional regulation and is expressed in sterile lemma primordia to prevent its development in the wild-type rice plant; it is abnormally enlarged like the lemma in the *g1* loss-of-function mutant (Yoshida et al. 2009). TAWAWA1 (TAW1), which is expressed in the inflorescence meristem and branch meristem, is also a transcription factor. TAW1 regulates the inflorescence architecture of rice by promoting inflorescence meristem activity and suppressing the phase change to spikelet meristem identity (Yoshida et al. 2013). For TRIANGULAR HULL1/BEAK LIKE SPIKELET1/ABNORMAL FLOWER AND DWARF1 (TH1/BLS1/ADF1), the mutants showed various pleiotropic defects containing abnormal floral organs, indicating that TH1/BLS1/ADF1 influences the cell size and number by negatively regulating the expression of genes related to cell proliferation and expansion (Li et al. 2012, Ma et al. 2013, Sato et al. 2014, Ren et al. 2016, Peng et al. 2017). In *Marchantia polymorpha*, LATERAL ORGAN SUPRESSOR 1 (LOS1) regulates meristem maintenance and lateral organ development by inhibiting cell division (Naramoto et al. 2019). Although not all ALOG proteins have the same molecular function despite the high sequence homology among ALOG domains (Naramoto et al. 2020), studies have shown that MpLOS1 can complement the phenotype of the rice *g1* mutant, indicating some conservation among their molecular functions (Naramoto et al. 2019). The suppressive effect of DAI on cell proliferation and elongation is consistent with the functions of previously reported ALOG proteins (Peng et al. 2017, Naramoto et al. 2019). Although the precise role of DAI as a transcription factor requires further investigation, our findings indicate functional conservation among ALOG proteins.

### Occurrence of DAI by gene duplication

Our finding that *DAI* may have originated from gene duplication of *DAI*^ori^ provides an interesting example of gain-of-function associated with altered morphologies of spikelet-related organs, similar to the case of another ALOG protein, G1, in rice. Consequently, its awn-specific expression during SS development (Fig. 6D) was assumed to confirm the inhibitory role of DAI. In this scenario, a duplicated gene in a region conferring suitable expression by the novel promoter may be critically important for gaining new functions. Supporting this assumption, we found that *DAI* acquired an awn-specific expression pattern that was distinct from that of *DAI*^ori^ (Fig. 6D). Additionally, the *DAI* genomic region containing its promoter can act as an awn inhibitor in rice, for which an inhibitor gene has not been identified. Together, these results suggest that the generation of dominant awn inhibitors depends on a rare event during the diversification of cereals and is thus species-specific.

The *DAI* insertion site was within a non-coding DNA region that is 15.48 and 10.26 kb away from upstream (Sobic.003G421201) and downstream (Sobic.003G421400) genes, respectively (Supplementary Fig. 2D). The upstream sequence of the insertion site in chromosome 3 may be sufficient to drive awn-specific expression; however, it remains unclear which promoter sequences are essential for the function of DAI. Because there is a putative long interspersed nucleotide element (LINE) upstream of the insertion site (Fig. 3A), whose epigenetic modifications lead to changes in the expression of nearby genes (Ong-Abdullah et al. 2015), it is possible that this LINE was involved in the establishment of *DAI* function. Thus, further analysis of the mechanism of transcriptional regulation of *DAI* is necessary.

In addition to DAI, the possible role of DAI^ori^ should be examined to improve the understanding of the ALOG protein family in sorghum. Because both proteins contain identical amino acids, DAI^ori^ may share its function with DAI in inhibiting cell proliferation and cell elongation in specific organs or tissues where *DAI*^ori^ is expressed. Although both *DAI* and *DAI*^ori^ transcripts accumulated in immature spikelets, their spatiotemporal expression patterns differed. Particularly, only *DAI*^ori^ transcripts accumulated substantially in the roots. Phylogenetic analysis showed that DAI/DAI^ori^ belongs to the G1L2 clade among other ALOG proteins (Fig. 4C). The precise roles of the proteins in the G1L2 clade are unknown, except for the results obtained in the present study on DAI. Considering its expression pattern (Fig. 6), DAI^ori^ may regulate root-related organs/tissues and spikelet-related organs/tissues other than the awn. Thus, further analysis of the functions of genes in the G1L2 clade, including *DAI*^ori^, is necessary.

### Additional two QTLs controlling awn length

Notably, two additional QTLs for awn length, *qAL7* and *qAL9*, were identified. Because information on such awn-related QTLs was limited in sorghum, we examined *qAL7* and *qAL9* further in our RIL population. Unlike *DAI*, these QTLs exerted additive effects on elongating awns, with NOG as the dominant allele. According to the QTL Atlas (Mace et al. 2019), no QTLs related to awn length have been reported around *qAL7* and *qAL9*. However, further investigation of the possible genes corresponding to these QTLs identified two possible candidate genes, *Dwarf1* (*Dw1*) and *Dwarf3* (*Dw3*), which are the major plant height-responsible genes in sorghum. *Dw1* encodes a positive regulator gene of brassinosteroid signaling and regulates the length of the internodes by controlling cell proliferation (Hilley et al. 2016, Yamaguchi et al. 2016, Hirano et al. 2017). *Dw3* encodes an ABCB1 auxin efflux transporter (Multani et al. 2003). Previously, we identified two dominant QTLs that control plant height using the same RILs, which corresponded to *Dw1* and *Dw3*. These two loci were shown to pleiotropically influence the size of spikelet-related organs, such as SSs, pedicels, anthers, and styles in the same RIL population (Takanashi et al. 2021). Considering that NOG has functional alleles for both genes (Kajiya-Kanegae et al. 2020), *qAL7* and *qAL9* detected in this study likely correspond to *Dw3* and *Dw1*, respectively.

### Conclusion

We identified the dominant awn-inhibiting gene in sorghum, which had not been evaluated for a century since its first mention (Vinall and Cron 1921). We showed that *DAI* encodes the ALOG protein, which appears to eliminate awn elongation by suppressing cell proliferation and elongation as a transcription factor. Experiments using sorghum germplasms suggested that *DAI* has been widely used in the process of modification or breeding of sorghum, perhaps because of its effectiveness and convenience. Further analysis of *DAI*, which differs from any other gene identified in other grass species, will provide insights into the mechanisms underlying awn regulation and generation of a novel gene via gene duplication.

## Materials and methods

### Plant materials

The RIL population, derived from a parental cross between BTx623 and the Japanese landrace NOG, was established as described previously (Kajiya-Kanegae et al. 2020). In total, 245 RILs were generated by recurrent selfing of progeny derived from a cross between BTx623 and NOG. A total of 289 accessions of sorghum germplasm was obtained from public bioresources (ICRISAT sorghum collection, NARO Genebank, and USDA); Supplementary Table 1 lists the accessions. Seeds of the rice cultivar Kasalath were kindly provided by Dr. Yasuo Nagato from the University of Tokyo. NILs were obtained starting from RIL205 (awnless), one of the heterozygous lines at the genotype of the marker associated with *qAL3/qAP3*, in the RILs of the F_6_ generation (Fig. 5A). We established NILs (F_9_ generation) through PCR genotyping using insertion-deletion polymorphisms for the recurrent selfed progeny of RIL205 (primers used for genotyping are listed in Supplementary Table 2).

### Genotype data

Genotype data of RILs were obtained using RAD-seq. The detailed methods and information for DNA extraction and library construction for RAD-seq were described in our previous report (Kobayashi et al. 2017). The RAD-seq reads of the 245 RILs from the F_6_ generation were mapped onto the reference genome *S. bicolor* v2.1 (Paterson et al. 2009) using the bwa mem algorithm (Li and Durbin 2010) with the default options, and then the marker positions were corrected using the information of the reference genome *S. bicolor* v3.1.1 (McCormick et al. 2018). SNPs were called using GATK (McKenna et al. 2010) and filtered with SAMtools (Li et al. 2009) under the following conditions: min-depth 5, max-depth 30, max-missing 0.8, minimum allele frequency 0.1, remove-indels, min-alleles 2, max-alleles 2, and minQ 20. For segregation distortion analysis, we used a chi-square test to calculate the deviation from the expected ratio (1:1) for each marker. All genotypes (BTx623/BTx623, NOG/NOG, and BTx623/NOG) were used to determine the allelic effects of QTLs in the F_6_ generation. To construct the genetic map for the RILs, only homozygous SNPs were selected, and those with missing rates greater than 20% and duplicated markers were removed. In total, 1,792 high-quality SNPs were selected to construct a linkage map for QTL analysis. The map covered a total length of 682.4 cM and had an average marker density of 0.4 cM.

Genotype data of the sorghum germplasms were also obtained using RAD-seq. The detailed methods and information for genotyping are described in our previous report (Yamazaki et al. 2020). SNPs with minor allele frequencies (1%) or missing rates greater than 80% were filtered out for the GWAS study. Imputation was conducted using Beagle 5.0 with default parameter settings (Browning et al. 2018). We obtained data for 50,678 SNP markers from 289 accessions.

### Cultivation

Field trials were performed in an open-air greenhouse at the experimental farm of the Institute of Plant Science and Resources (IPSR) at Okayama University (latitude: 34°35′31” N, longitude: 133°46′7″ E) from June to September 2014 for the F_6_ generation of the RILs. Plants were germinated in small trays filled with vermiculite for approximately 10–14 days and then transplanted to 30-cm diameter pots filled with soil from the IPSR field (n = 1 per line).

Field trials were performed in a green field at the University of Tokyo (latitude: 35°42’58.5” N, longitude: 139°45’44.5’’ E) from May to October 2015, 2019, and 2020. Plants were germinated in 200-cell trays filled with synthetic culture soil (BONSOL 1; Sumitomo Chemical Co., Ltd., Tokyo, Japan) for two weeks, and then transplanted into the field (n = 2 per line). The space between ridges was 60 cm, with a 15 cm distance between individuals in the same ridge. The field was fertilized using N:P:K =10:10:10 (kg ha^-1^) fertilizer before the trials each year. Cultivation and phenotyping of awn-related traits for RILs were conducted in 2015 using the F_7_ generation, and those for the germplasm were conducted in 2015, 2019, and 2020.

Transgenic sorghum and rice plants were grown and maintained in a biohazard greenhouse at 30 °C (day) and 25 °C (night) under natural light conditions.

### Phenotyping for QTL analysis and GWAS

To standardize the developmental stages of SSs among the RILs and germplasm, we sampled SSs at the just-before-flowering stage in each line, except for in the F_6_ generation of the RILs cultivated in Okayama, which were sampled during the grain-filling stage. In the inflorescence that began flowering at the top, we defined SSs at the just-before-flowering stage (SSs that would flower within a day) as those attached to the primary branch immediately below the flowered primary branch. For each line, we picked awns from five SSs sampled from the top, middle, and bottom of the panicle, mounted them on a paper board, and photographed them using a digital camera (OM-D E-M5; Olympus, Tokyo, Japan). The AL was measured using the ImageJ 1.49v software (Schneider et al. 2012) (NIH, Bethesda, MD, USA). We also scored awn presence values on a two-step scale with awnless lines as 0 and awned lines as 1. For all awn-related traits, the average value of five independent awns was used as the phenotypic value for an individual plant.

### QTL analysis

We performed QTL analysis for awn-related traits in the RILs (AL, AP, and AL only in awned lines). Genotype probabilities were calculated using calc.genoprobability function with a step size of 1 cM and an assumed genotyping error probability of 0.05, using the Kosambi map function (Kosambi 1943) as implemented in the R (R Core Team 2021)/qtl package (Broman et al. 2003). QTL analysis was performed using the composite interval mapping function of the R/qtl package with the Haley–Knott regression method (Haley and Knott 1992). Linkage analysis was performed using the R/qtl package in R, version 4.1.0. The logarithm of odds (LOD) significance threshold for detecting QTLs was calculated by performing 1,000 iterations using the R/qtl permutation test. Confidence intervals for the QTLs were estimated based on the 2.0-LOD support interval, and the nearest flanking markers outside the boundary of each confidence interval were defined as both ends of the marker intervals. The additive effect and percentage of phenotypic variance explained by each QTL were obtained using the fitqtl function of the R/qtl package. Box plots were created using the R/ggplot2 package (Wickham 2016).

### GWAS

We performed GWAS of awn-related traits (AL and AP) in the germplasm. GWAS was performed using a linear mixed model implemented by the “association.test” function in the gaston package ver. 1.5.7 in R (Perdry et al. 2020). A genetic relationship matrix specifying a random additive effect was computed using the ‘A.mat’ function in the rrBLUP package (Endelman 2011). The P-values of the marker-trait associations were calculated using the Wald test. The genome-wide significance threshold was obtained based on the Benjamini–Hochberg false discovery rate (Benjamini and Hochberg 1995) at 0.1% level. Linkage disequilibrium heatmaps surrounding the peaks in the GWAS were constructed using the “LD” function of the gaston package. To determine the geographic distribution of alleles at the peak SNP in GWAS, a world map layout was generated using the ggplot2 package in R. Passport data of accessions were obtained from the U.S. National Plant Germplasm System (https://npgsweb.ars-grin.gov/gringlobal/search), NARO Genebank (https://www.gene.affrc.go.jp/databases-core_collections_ws.php), and ICRISAT sorghum collection (http://www.icrisat.org/what-we-do/crops/sorghum/Project1/Sorghum_minicore.htm).

### Genotyping of 5.46 kb fragment containing DAI for BTx623, NOG, and selected germplasm

Healthy young leaves from each line were collected and genomic DNA was extracted using a Maxwell RSC Plant DNA Kit (Promega, Madison, WI, USA). The presence or absence of the 5.46-kb fragment containing *DAI* was confirmed by PCR (primers 1 and 2), and sequence information was determined by direct Sanger sequencing using the same primers.

### Plasmid construction

To transiently express GFP and DAI-GFP, we constructed the corresponding constructs using Gateway cloning technology (Invitrogen, Carlsbad, CA, USA). The CDS of *DAI* without a stop codon was amplified by PCR using BTx623 DNA as a template (primers 3 and 4), and the amplified DNA fragment and an annealed linker oligo (for GFP: primers 5 and 6) were introduced into the pENTR/D-TOPO vector (Invitrogen; sequences were confirmed by Sanger sequencing using primers 7 and 8) and then subcloned into a pH7FWG2 vector (Karimi et al. 2002) using LR clonase (Invitrogen) to obtain 35Spro::DAI-GFP and 35Spro::GFP constructs.

For the sorghum and Kasalath transgenic lines, the constructs were prepared using In-Fusion HD cloning technology (Clontech, Mountain View, CA, USA). For *DAI*pro::*DAI*, the pBUH3 vector (Nigorikawa et al. 2012) was linearized using PCR (primers 9 and 10). The DNA fragment containing the *DAI* native promoter and CDS of *DAI* with a stop codon were amplified by PCR using the *SbDAI* plasmid as a template (primers 11 and 12), and the PCR-amplified DNA fragment was cloned into linearized pBUH3 via In-Fusion HD cloning. The sequence of the *DAI*pro::*DAI* plasmid was confirmed by Sanger sequencing using primers 13–18. For *SbDAI*, the DNA fragment containing the *DAI* native promoter and CDS of *DAI* with a stop codon were amplified by nested PCR using BTx623 DNA as a template (primers 25–28). For gNOG, the DNA fragment was amplified by nested PCR using NOG DNA as a template (primers 25, 2, 27, and 29). Amplified *DAI*pro::*DAI* and gNOG fragments were cloned into the *Nco*I/*Sac*I-digested pHGWFS7 vector (Karimi et al. 2002) via In-Fusion HD to generate the *SbDAI* and gNOG plasmids, respectively. The sequences of these plasmids were confirmed by Sanger sequencing using primers 16 and 30–33.

For standard plasmids for qRT-PCR, the DNA fragment of the 3’ untranslated region of *DAI* that matched perfectly for *DAI* and *DAI*^ori^ and the DNA fragment of *SbTFIIE* (transcription initiation factor IIE: *Sobic. 006G209500*) were amplified by PCR using BTx623 DNA as a template (primers 70–73) and cloned into the pENTR/D-TOPO vector to obtain pENTR_Comm and pENTR_*TFIIE*, respectively.

### Particle bombardment

The GFP and DAI-GFP plasmids were transiently introduced into onion epidermal cells using a helium-driven particle accelerator (PDS/1000; Bio-Rad Laboratories, Hercules, CA, USA). The particle bombardment experiments were performed as described by Arimura et al. (2004). Before microscopic observation, onion epidermal cells were treated with distilled water containing 2 mg L^-1^ 4’,6-diamidino-2-phenylindole dihydrochloride to stain the nuclei.

### Microscopic observations

Enlarged images of the secondary branches and spikelets of sorghum and Kasalath were captured under a stereomicroscope (M125; Leica, Wetzlar, Germany) using a CCD camera (MC170 HD; Leica). Onion epidermal cells expressing GFP and DAI-GFP under control of the *35S* promoter and calcofluor white-stained dissected awns were observed using a Leica STELLARIS5 confocal microscope (Leica) with an HC PL APO CS2 20× DRY objective lens (numerical aperture = 0.75), HC PL APO CS2 63× OIL objective lens (numerical aperture = 1.40), and HyD detectors. The Leica STELLARIS5 microscope was controlled using the LAS X software (Leica); 4’,6-diamidino-2-phenylindole dihydrochloride/calcofluor white and GFP were excited using 405 nm diode laser and 495 nm tuned white light lasers, respectively.

### Agrobacterium-mediated transformation

*Agrobacterium*-mediated sorghum transformation of RIL232 (one of the lines in our RILs, awned) was performed according to Wu et al. (2014), with several modifications. Immature embryos isolated from the grains at 15 days after pollination were pre-cultured in DBC medium at 25 °C in the dark. Pre-cultured immature embryos were collected in the infection medium, pre-treated at 43 °C for 15 min, and centrifuged (20,000 ×*g* for 10 min). Pre-treated immature embryos were inoculated with *Agrobacterium tumefaciens* strain EHA105 harboring the *DAI*pro::*DAI* vector and co-cultured at 25 °C for 5 days in the dark. After 5 days of resting culture, calli were selected on medium containing 15–20 mg L^-1^ hygromycin, and the resistant calli were transferred to a regeneration medium containing 10 mg L^-1^ hygromycin. Regenerated shoots were cultured on rooting medium until roots were produced. We obtained several plantlets; however, only one was confirmed to be a transformant by PCR. Segregation analyses of T_1_ plants derived from the self-pollinated T_0_ plant by PCR using primers 19 and 20 revealed that the T_0_ plant was heterozygous for a single copy of the transgene (segregation ratio of approximately 3:1). We confirmed the number of transgenes in segregated T_1_ plants by qPCR using primers 21–24 and selected five individuals each as homozygous, heterozygous, and azygous for AL measurement. For each line, we picked awns from 30 SSs sampled from the middle portion of the panicle at the just-before-flowering stage and measured the AL as described above.

The constructed gNOG and *SbDAI* vectors were transformed into calli induced from the scutellum of wild-type (cv. Kasalath, awned) using *A. tumefaciens* (strain EHA105), as described by Toki et al. (2006). Transgenic plants were selected on medium containing 50 mg L^-1^ hygromycin. Hygromycin-resistant plants (T_0_) were transplanted into the soil and grown at 30 °C (day) and 25 °C (night). Genotyping of T_0_ plants was performed by PCR using primers 19, 34, and 35, and three independent transgenic lines were obtained for each construct. T_1_ plants derived from the self-pollinated T_0_ plants were genotyped by PCR, and individuals with or without transgene(s) were selected for AL measurement. For each line, we sampled the panicles before flowering and measured the AL of apical spikelets on each primary and secondary branch (approximately 100 spikelets).

### Synteny analysis

Synteny analysis was performed using an online SynMap2 software (Haug-Baltzell et al. 2017). Datasets were selected for dot plots as follows: *Sorghum bicolor* (id331), *Oryza sativa* (id3), *Brachypodium distachyon* (id39836), and *Zea mays* (id38215). Only the results of the chromosomes corresponding to *DAI* (Sb_Chr03) and *DAI*^ori^ (Sb_Chr10) were extracted from the output files of SynMap2, and dot plots were drawn using ggplot2/R. Enlarged images around *DAI* or *DAI*^ori^ in Figure 5B are based on the reference genomes of *S. bicolor* (v3.1.1), *O. sativa* (v7.0), *B. distachyon* (v3.1), and *Z. mays* (RefGen_V4) in Phytozome 13 (Goodstein et al. 2012).

### Phylogenetic analysis

The amino acid sequence of Sobic.003G421300/DAI was used as the query sequence for a BLASTP search against the protein databases of *S. bicolor* (v3.1.1), *O. sativa* (v7.0), *B. distachyon* (v3.1), and *Z. mays* (RefGen_V4) using Phytozome 13. Fifty matches with expected values less than e^-40^ were selected, and a phylogenetic tree was constructed using ClustalW alignment and maximum likelihood methods based on the JTT matrix-based model (Jones et al. 1992) using the default settings in the MEGA 7 package (Kumar et al. 2016). The following protein sequences from the Phytozome database were used for phylogenetic analysis. *S. bicolor:* Sobic.001G219400, Sobic.002G029700, Sobic.003G342600, Sobic.003G421300 (DAI), Sobic.004G052800, Sobic.004G219600, Sobic.004G338400, Sobic.005G094266, Sobic.006G147500, Sobic.009G102600, Sobic.009G170100, and Sobic.010G225100 (DAI^ori^); *O. sativa:* LOC_Os01g61310 (G1L7), LOC_Os02g07030 (G1L1), LOC_Os02g41460 (G1L3), LOC_Os02g56610 (G1L6/TH1/BLS1/ADF1), LOC_Os04g43580 (G1L4), LOC_Os05g28040 (G1L9), LOC_Os05g39500 (G1L8), LOC_Os06g46030 (G1L2), LOC_Os07g04670 (G1), and LOC_Os10g33780 (G1L5/TAW1), *B. distachyon:* Bradi1g32445, Bradi1g58070, Bradi2g22561, Bradi2g29110, Bradi2g53880, Bradi3g04960, Bradi3g28800, Bradi3g49140, Bradi3g54800, and Bradi5g15700; and *Z. mays*: Zm00001d003057, Zm00001d010507, Zm00001d010596, Zm00001d012473, Zm00001d014032, Zm00001d015342, Zm00001d017263, Zm00001d018433, Zm00001d018775, Zm00001d025863, Zm00001d032696, Zm00001d036617, Zm00001d038016, Zm00001d038442, Zm00001d043011, Zm00001d046998, Zm00001d051143, and Zm00001d051966.

### Observation of dissected awns of immature SSs in NILs

Immature SSs at stages 5 and 7 in both the NILs were sampled before heading and dissected under a stereomicroscope. Dissected awns were cleared using ClearSee (Kurihara et al. 2015) for 2 days. The cell walls of cleared samples were stained with 1 g L^-1^ calcofluor white (fluorescent brightener 28, Sigma, St. Louis, MO, USA) in ClearSee for 1 h and washed with ClearSee for 1 h. The stained samples were observed using confocal laser scanning microscopy. The DALs, numbers of adaxial epidermal cells, and lengths of adaxial epidermal cells in the three regions (bottom, middle, and top; see Fig. 5E) were measured using the ImageJ 1.49v software. Ten dissected awns of both NILs were measured for each stage.

### Quantitative RT-PCR

Fully expanded 5^th^ leaves and lateral roots were collected from 2-week-old NIL_B plants. Immature panicles were sampled from NIL_B plants before heading, and primary branches at each stage were collected under a stereomicroscope. Total RNAs for quantitative RT-PCR were extracted from each stage of immature primary branches, leaves, and roots using a Maxwell RSC Plant RNA kit (Promega) according to the manufacturer’s instructions. We prepared three biological replicates for immature primary branches, leaves, and roots. As the SSs at earlier stages were too small to isolate all the intact components, we used the SSs in NIL_B at stage 8 for analysis. The SSs at stage 8 were carefully dissected under a stereomicroscope, and each component from the 100 SSs was collected and pooled. Total RNAs were extracted from the SS components using an Arcturus PicoPure RNA isolation kit (Thermo Fisher Scientific, Waltham, MA, USA).

First-strand cDNA was synthesized from 1 μg of each RNA sample in a 21 μL reaction mixture volume using SuperScript IV Reverse Transcriptase (Thermo Fisher Scientific). The transcript levels were measured using a StepOnePlus real-time PCR system (Applied Biosystems, Foster City, CA, USA) and PowerUp SYBR Green Master Mix (Thermo Fisher Scientific). *DAI* and *DAI*^ori^ have high sequence homology (see Supplementary Fig. 3), making it difficult to design primers that specifically amplify each gene. After repeated examinations, we designed specific primers for *DAI* but not for *DAI*^ori^. Therefore, we calculated the quantitative value of *DAI*^ori^ by subtracting that of *DAI* from the total quantitative value of *DAI* and *DAI*^ori^ estimated using the primer set amplifying *DAI* and *DAI^ori^* together. The plasmids containing each target sequence were quantified and used to generate standard curves for absolute quantification. Transcript levels were normalized to the level of the internal control gene *SbTFIIE*. The plasmids and primers used to draw the standard curves were *SbDAI* (in pHGWFS7) for *DAI:* primers 76 and 77, pENTR_Comm for *DAI* + *DAI*^ori^: primers 71 and 74, and pENTR_*TFIIE* for *SbTFIIE:* primers 72 and 75. Data are expressed as the average of three biological replicates in immature primary branches, leaves, and roots (Fig. 6C) and three technical replicates in components of the SS (Fig. 6D).

## Data availability

The sequence data reported in this paper have been submitted to the DNA Data Bank of Japan Sequence Read Archive under accession number DRA008159.

## Author contributions

Material preparation, field experiments, and data analysis in QTL and GWAS were performed by H.T., H.K.-K., W.S., M.I., with supervision from M.K., K.Y., H.I., and N.T.; NIL was established by W.S and analyzed by H.T. Sorghum transformation was performed by A.N., J.Y., and H.T. with supervision from N.T.; Rice transformation was performed by H.T.; final data were prepared for publication by H.T., H. K.-K., and W.S.; and the manuscript was written by H.T. and W.S., on behalf of all the authors.

## Funding

This work was partly supported by the Core Research for Evolutional Science and Technology (CREST) from the Japan Science and Technology Agency (to N.T. and W.S.) and KAKENHI grants (17H01457 to N.T., 18K19343 and 21H02597 to W.S., 18K05570 to H.T.) from the Japanese Society for the Promotion of Science (JSPS). We also thank the Oohara Foundation for the financial support of our research group.

## Supplementary Data

Supplementary data are available at PCP online.

## Disclosures

The authors have no conflicts of interest to declare.

## Acknowledgments

We thank Tsuneaki Takami, Zihuan Jing, Fiona Wacera, Norikazu Ohnishi, Everlyne Omollo, and Rie Hijiya (IPSR, Okayama University) for their help with cultivation at the IPSR, Kurashiki, Okayama. We are also grateful to the members of the Laboratory of Plant Molecular Genetics and the Laboratory of Biometry and Bioinformatics at the University of Tokyo for their help with cultivation in Tokyo.

---

**Supplementary Figure 1.**
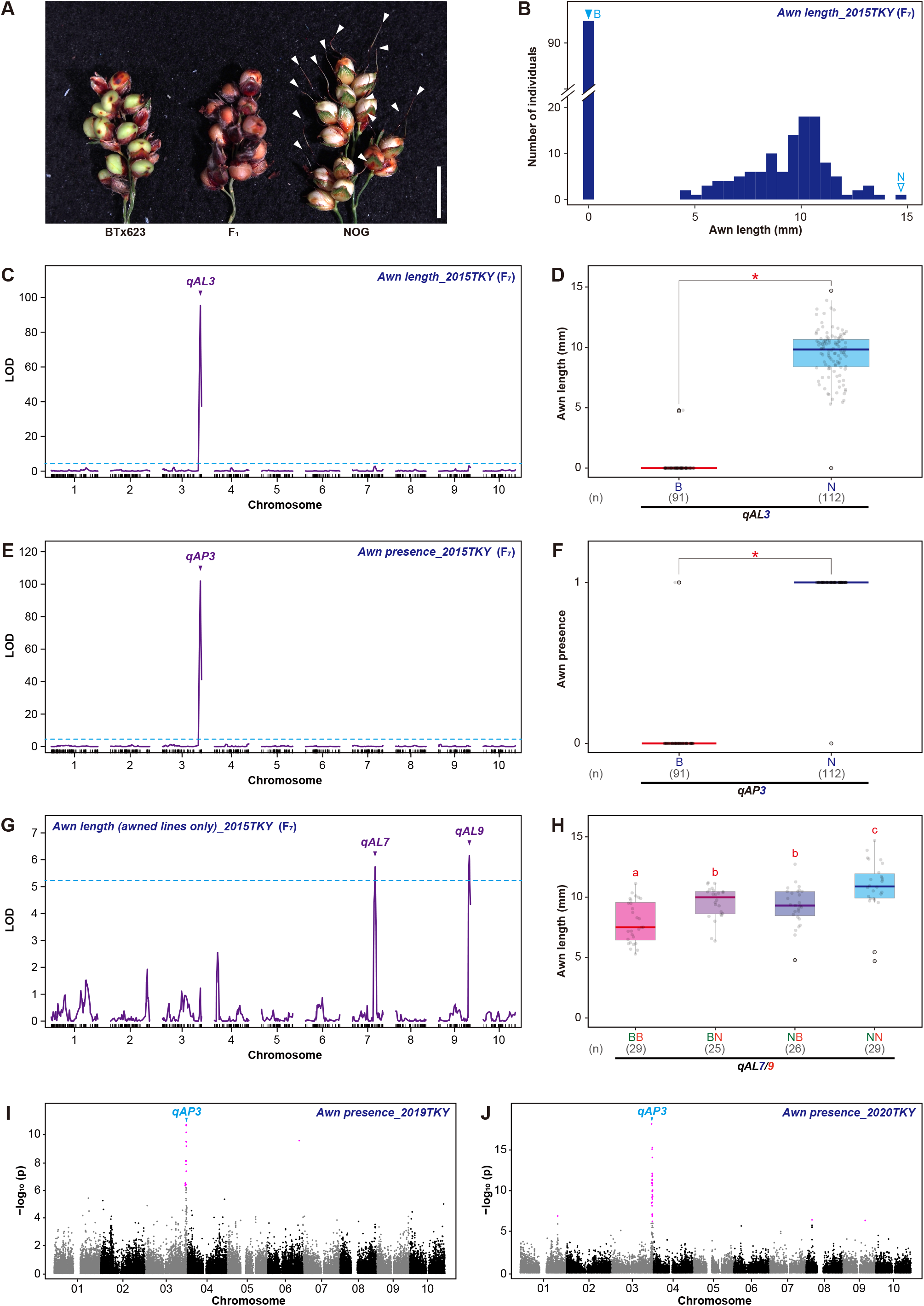
Reproducibility check of quantitative trait locus (QTL) analysis and genome-wide association study using lines cultivated in different years [2015 at Tokyo for recombinant inbred lines (RILs) (F_7_ generation) and 2019 and 2020 at Tokyo for germplasm] from those in Fig. 1 and 2. (A) Awn phenotypes of F_1_ and its parents. Arrowheads indicate awn tips. (B) Frequency distribution of awn length (AL) in the RIL population (F_7_ generation) cultivated at Tokyo in 2015. A filled arrowhead and an open arrowhead indicate phenotypic values for BTx623 and NOG, respectively. Results of QTL analysis for the awn-related traits (C, E, G) and their allelic effects (D, F, H). (C, D) AL of all lines, (E, F) awn presence (AP), (G, H) AL of awned lines only. (C, E, G) Logarithm of odds (LOD) profiles obtained from composite interval mapping (CIM). Horizontal dotted lines represent a threshold of the 1,000× permutation test (*P* < 0.05). (D, F, H) Contributions of single-nucleotide polymorphism (SNP) genotypes of the nearest marker for significant QTLs. Box plots show the effects of the nearest marker genotypes for each QTL or allelic combinations of QTLs. B, BTx623-type homogeneous allele; N, NOG-type homogeneous allele. The number of lines corresponding to each genotype is shown in parentheses. Asterisks indicate significant differences between genotypes (Welch’ s *t*-test, *P* < 0.001). Different letters denote significant differences according to the Tukey–Kramer test (*P* < 0.05). (I, J) Manhattan plots for AP in 2019 and 2020. Dots in magenta indicate the SNPs significantly associated with traits (Benjamini–Hochberg-corrected significance, *P* < 0.001). Scale bars = 1 cm in (A).

**Supplementary Figure 2.**
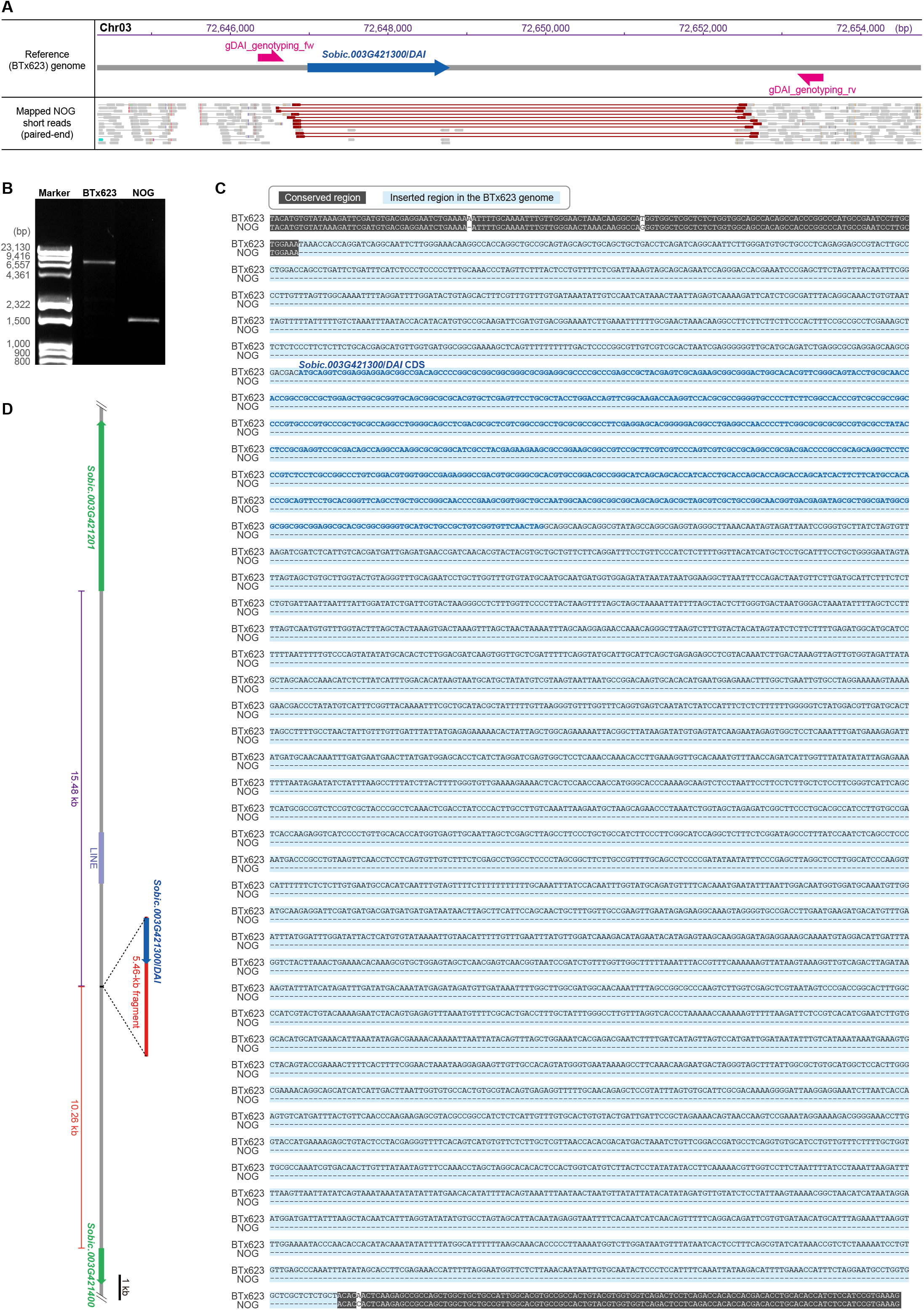
Difference between genomic structures of BTx623 and NOG around *DAI*. (A) Re-sequencing data of NOG in the region around *DAI* visualized using Integrative Genomics Viewer. Pentagonal boxes represent each short read, and paired reads are connected to each other by lines. The paired reads shown in red indicate that the size of inserts were unexpectedly large, i.e., the NOG genome did not contain regions between them. (B) Result of genotyping of BTx623 and NOG by PCR using primers indicated in (A). (C) Details of the genomic structure of NOG and BTx623 around *DAI* as revealed by Sanger sequencing of PCR products in (B). (D) Insertion position of the 5.46-kb fragment and its relationship to nearby genes.

**Supplementary Figure 3.**
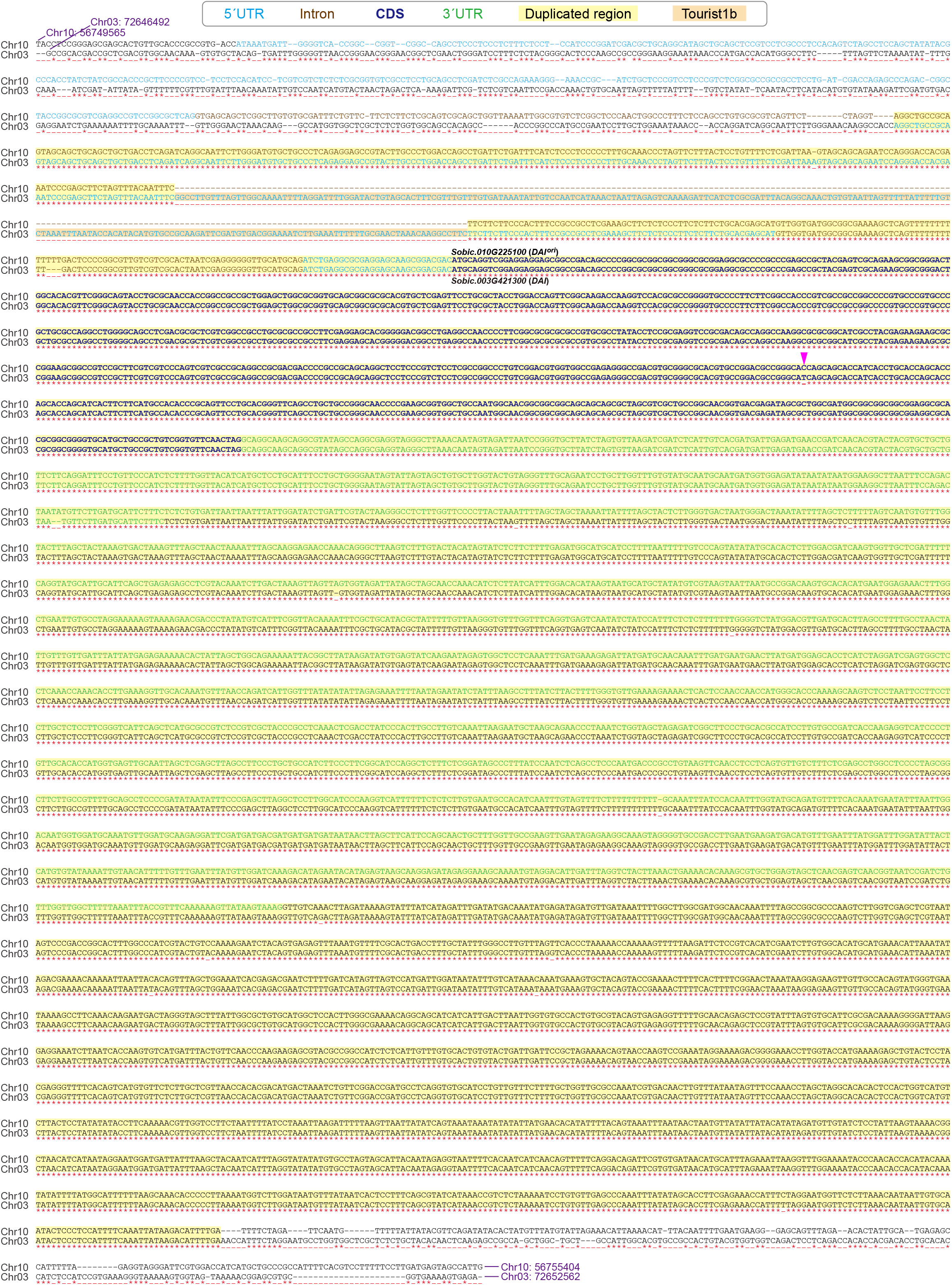
Details and relation of the genomic structure of the region around *DAI*^ori^ (Chr10) and *DAI* (Chr03) in the BTx623 genome. Arrowhead in magenta indicates the only base substitution (synonymous substitution) that exists between the coding sequence of *DAI*^ori^ and that of *DAI*.

## Notes

### Competing Interest Statement

The authors have declared no competing interest.

### Summary of Updates

In the revised manuscript, additional experiments were included to address i) the suppressing mechanism of awn elongation mediated by DAI, and ii) the expression level of DAI and its ancestor DAIori. These additional results allowed us to conclude that DAI inhibits awn elongation by suppressing both cell proliferation and cell elongation, and that DAI acquired awn-specific expression pattern distinctive to DAIori. Title is accordingly modified. In addition, we re-organized the text so as to have more focus on the above findings.

